# Hematopoietic Stem and Progenitor Cell Aging is Initiated at Middle Age Through Decline in Local Insulin-Like Growth Factor 1 (IGF1)

**DOI:** 10.1101/2020.07.11.198846

**Authors:** Kira Young, Elizabeth Eudy, Rebecca Bell, Matthew Loberg, Tim Stearns, Lars Velten, Simon Haas, Jennifer J. Trowbridge

## Abstract

Hematopoietic stem cells (HSCs) are responsible for lifelong maintenance and regeneration of the blood system. With aging, loss of HSC function is a major contributor to decline in overall hematopoietic function, leading to increased rate of infection, poor vaccination response, clonal hematopoiesis, and increased risk of hematologic malignancies. While cellular and molecular hallmarks of HSC aging have been defined^1–3^, the lack of understanding of the nature and timing of the initiating events that cause HSC aging is a barrier to achieving the goal of extending healthy hematopoietic function into older age. Here we discover that hallmarks of HSC aging and myeloid-biased hematopoiesis accumulate by middle age in mice, and that the bone marrow (BM) microenvironment at middle age induces and is indispensable for hematopoietic aging phenotypes. Using unbiased transcriptome-based approaches, we identify decreased production of IGF1 by cells in the middle-aged BM microenvironment as a factor causing hematopoietic stem and progenitor cell aging and show that direct stimulation with IGF1 rescues hallmarks of hematopoietic aging. Declining IGF1 in the BM microenvironment at middle age represents a compelling target for intervention using prophylactic therapies to effectively extend healthspan and to prevent functional decline during aging.

## Main

Hallmarks of HSC and hematopoietic system aging in mice and humans have been largely defined by the binary comparison between young and old individuals^1,2,4–8^, limiting the understanding of mechanisms causing aging versus alterations that are consequences of aging. To identify initiating mechanisms of aging, we employed a cross-sectional analysis of C57BL/6J mice. In female mice, we found that by 9-12 months of age (corresponding to ~36-45 years of age in humans^9^) many hallmarks of hematopoietic and HSC aging were observed, including increased frequency of myeloid cells relative to lymphoid cells in the blood^7^ (Fig. 1A, Extended Data Fig. 1A), increased frequency of phenotypic long-term HSCs (LT-HSCs)^10,11^ (Fig. 1B, Extended Data Fig. 1B, gating strategy shown in Extended Data Fig. 1C), increased frequency of LT-HSCs with γH2AX foci^12^ (Fig. 1C), and loss of polarity of CDC42 and tubulin^13^ (Fig. 1D). In male mice, we also observed increased frequency of myeloid cells relative to lymphoid cells in the blood, but this occurred beyond 9-12 months of age (Extended Data Fig. 1D), indicating potentially distinct sex-specific dynamics^14^.

**Fig. 1:**
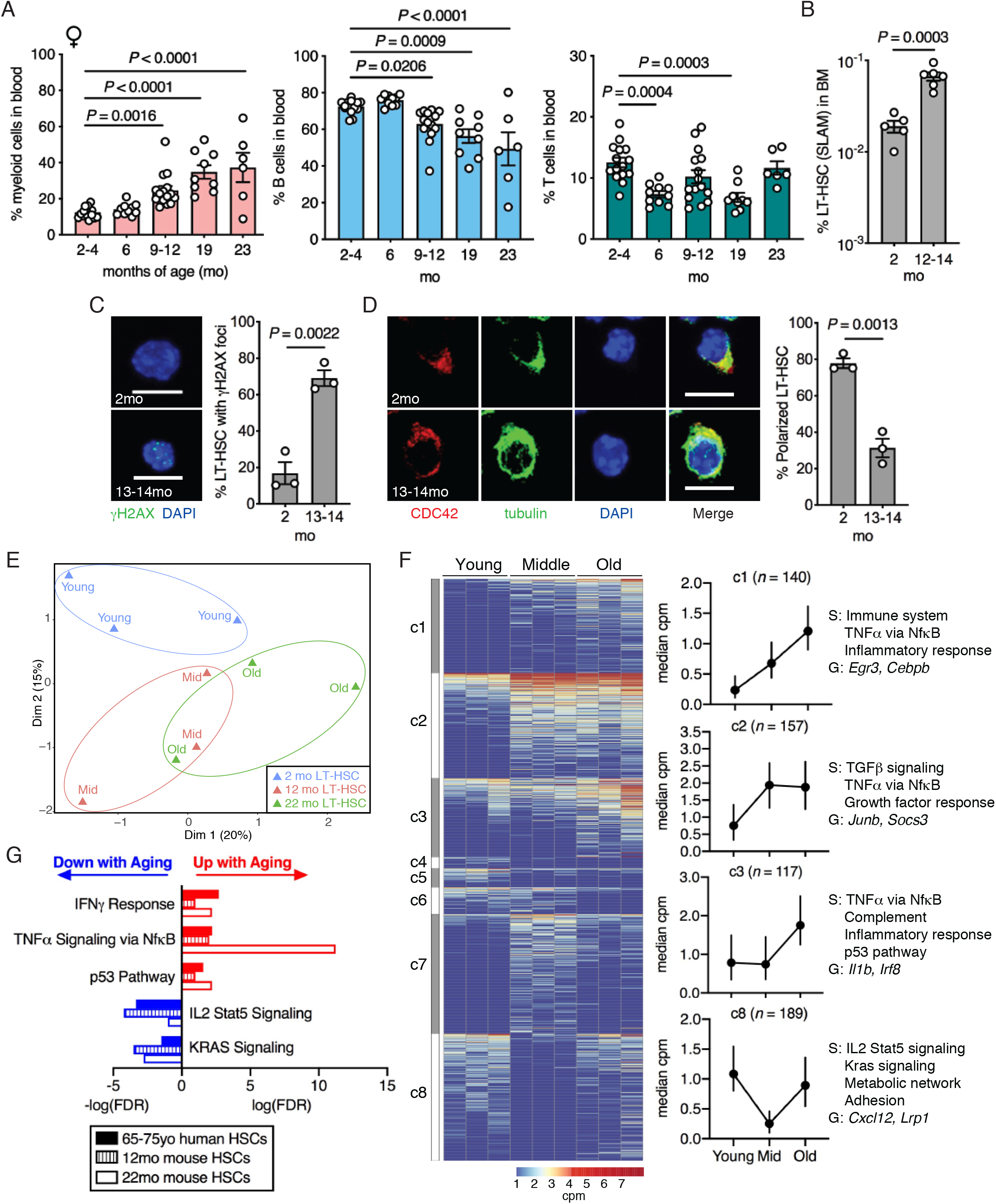
Hallmarks of LT-HSC aging occur by middle age in mice. (**a**) Frequency of myeloid cells (CD11b+), B cells (B220+) and T cells (CD3+) within the blood of female mice at 2-4mo (*n =* 15), 6mo (*n =* 10), 9-12mo (*n =* 15), 19mo (*n =* 9) and 23mo (*n =* 6). (**b**) Frequency of LT-HSCs (defined using SLAM markers) in whole BM of mice at 2mo (*n =* 5) and 12-14mo (*n =* 6). (**c**) Left: Representative images of LT-HSCs from 2mo and 13-14mo mice stained with γH2AX and DAPI. Scale bar, 10 μm. Right: Quantification of the percentage of LT-HSCs with γH2AX foci from 2mo (*n =* 3) and 13-14mo (*n =* 3). (**d**) Left: Representative images of LT-HSCs from 2mo and 13-14mo mice stained with CDC42, tubulin, DAPI, and overlay. Scale bar, 10 μm. Right: Quantification of the percentage of LT-HSCs with polarized CDC42 and tubulin from 2mo (*n =* 3) and 13-14mo (*n =* 3). (**e**) Multidimensional scaling (MDS) plot of the top 500 differentially expressed genes (DEGs) identified by RNA-seq data from *de novo* isolated young (2mo) (*n =* 3), middle-aged (mid; 12mo) (*n =* 3) and old (22mo) (*n =* 3) mice. (**f**) Left: Heatmap of expression of all DEGs between young and middle-aged or young and old LT-HSCs. Denoted clusters (c1-c8) were derived based on similarity of expression patterns. Right: Expression graphs of selected gene clusters showing trends from young to middle-aged to old LT-HSCs (dots represent median, bars represent interquartile range, *n* represents number of genes in each cluster), along with enrichment of hallmark gene signatures (S), and selected genes (G) from each cluster. (**g**) Bar plot representation of hallmark gene sets enriched in DEGs from aged human HSCs^2^, middle-aged (12mo) mouse LT-HSCs and old (22mo) mouse LT-HSCs versus their respective young controls. (**a**-**d**) Dots represent individual mice and bars are mean ± SEM. All *n* values refer to the number of mice used. P-values were generated for (**a**) by one-way ANOVA with Holm-Sidak’s multiple comparisons test, (**b**-**d**) by unpaired, two-tailed *t* test.

To further investigate the extent to which middle-aged LT-HSCs resemble old LT-HSCs, we performed RNA sequencing from young (2mo), middle-aged (12mo) and old (22mo) *de novo* isolated LT-HSCs. Multidimensional scaling of this data revealed that there is modest overlap between middle-aged and old LT-HSCs, and that these are both distinct from the young LT-HSC signature (Fig. 1E). This indicates that middle-aged and old LT-HSCs are more transcriptionally similar to each other than young LT-HSCs but also that middle age may represent a distinct transcriptional state. To assess progressive changes during aging, we derived clusters from all significantly differentially expressed genes (FDR < 0.05, fold change (FC) > 1.5) (Fig. 1F), modeling distinct patterns of expression from young to middle-aged to old LT-HSCs. Clusters 1, 2 and 3 (c1 to c3) represented progressively increased expression, peak expression at middle-age that is sustained in old, or peak expression exclusively in old, respectively. These clusters were enriched for similar gene signatures, including immune and inflammatory response, and TNFα signaling. This suggests that while distinct inflammation-related gene modules are engaged in middle-aged versus old LT-HSCs, the overall inflammatory response is likely progressive and cumulative with age. Cluster 8 (c8), representing genes decreased specifically at middle age, was enriched for gene signatures of IL2-Stat5 signaling, Kras signaling, metabolic network and adhesion, suggesting that these pathways may define a distinct, “middle-aged” transcriptional state. We next examined how middle-aged and old LT-HSC expression signatures relate to previously defined myeloid-biased and lymphoid-biased LT-HSCs signatures^15^. We observed enrichment of myeloid-biased LT-HSC and depletion of lymphoid-biased LT-HSC signatures in both middle-aged and old LT-HSCs (Extended Data Fig. 1E). In addition, many pathways and processes altered in aged human HSCs (65-75 years old)^2^ were enriched in middle-aged and/or old mouse LT-HSCs, including increased IFNγ response, TNFα signaling, and the p53 pathway, and decreased IL2-Stat5 and Kras signaling (Fig. 1G). Despite significant overlap at the pathway level, examination at the gene level determined that only five genes overlap between aged human HSCs, middle-aged mouse LT-HSCs, and old mouse LT-HSCs, compared to their respective young controls, including *Egr1, Socs3, Selp* and *Fzd1* (Extended Data Fig. 1F; Supplementary Table 1). While this is greater overlap at the gene level than would be expected by chance (*P* < 2.26×10^−6^, Fisher’s Exact Test), these results suggest that overlap at the pathway level is driven to a greater degree by distinct genes in aging mouse HSCs and aging human HSCs that regulate common pathways. Together, these findings support that many molecular hallmarks of HSC aging are conserved at the pathway level between mice and humans, and occur as early as middle age in mice. These findings, and the unique transcriptional state of LT-HSCs at middle age, define a novel platform and molecular signature to identify and evaluate efficacy of interventions to extend hematopoietic and immune system healthspan.

Whether HSC-intrinsic or extrinsic changes, or a combination of both factors, cause hematopoietic aging is a matter of ongoing debate^3,16–1^. To evaluate the relative contribution of HSC-intrinsic versus extrinsic changes at middle age, we performed reciprocal transplantation of young, purified LT-HSCs into middle-aged mice (9 months of age) or middle-aged LT-HSCs (12 months of age) into young mice. Purified LT-HSC transplantation into lethally irradiated recipients requires infusion of mature cells to support hematopoiesis through irradiation recovery. As the age of these “support” cells may also influence LT-HSCs, we tested this as a variable in our experimental design. We analyzed the experiment at 24 weeks post-transplant, a time point at which mature hematopoietic lineage composition in both young and middle-aged recipients had returned to steady-state levels, indistinguishable from native non-transplant, age-matched mice (Extended Data Fig. 2A, B). We found that transplant of young LT-HSCs into middle-aged (9 months) recipient mice (YMM; Fig. 2A) increased myeloid cell production and decreased B and T lymphocyte production without altering overall engraftment level (Fig. 2B, Extended Data Fig. 2C), resembling the transplant of middle-aged LT-HSCs into middle-aged recipient mice controls. This phenotype was not reproduced upon transferring middle-aged “support” cells alone (YMY), indicating that the age of the recipient mouse was causative. In the reciprocal experiment, transplant of middle-aged LT-HSCs (12 months) into young recipient mice (MYY; Fig. 2C) restored myeloid and B cell production as well as engraftment level to that observed in young controls (Fig. 2D, Extended Data Fig. 2D). This phenotype was partially recapitulated in the MMY condition, suggesting that the age of support cells can modify engraftment in young recipient mice.

**Fig. 2:**
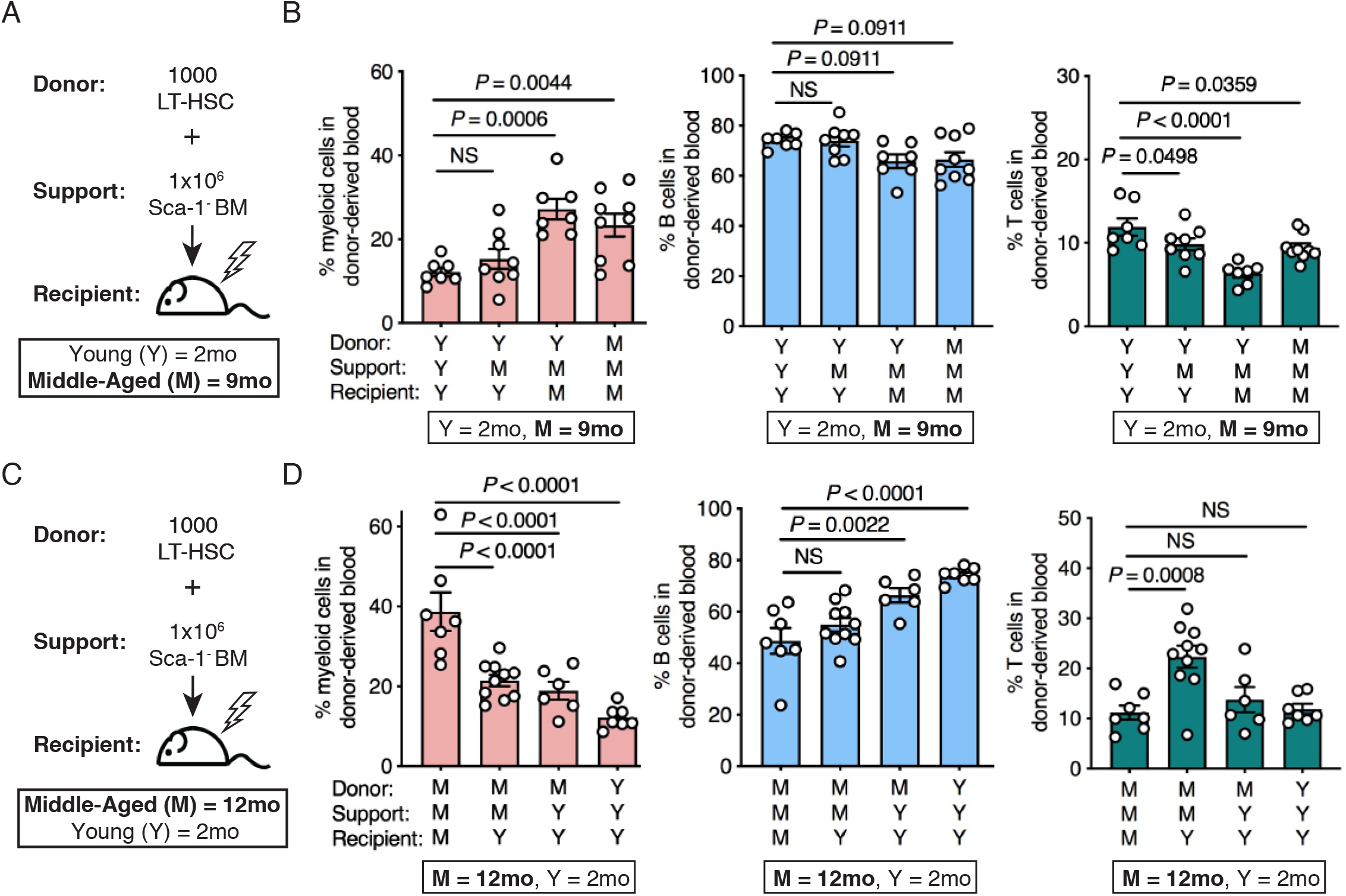
The middle-aged BM microenvironment is sufficient to cause myeloid-biased hematopoiesis. (**a**) Schematic of transplantation design using purified LT-HSCs (CD45.2+) plus support cells (CD45.1+ Sca-1-) into recipient mice (CD45.1+), where each component can include young (Y; 2mo) or middle-aged (M; 9mo) sources. (**b**) Frequency of donor-derived (CD45.2+) myeloid (CD11b+) cells, B cells (B220+) and T cells (CD3+) in the blood at 24wks post-transplant (*n* = 7, 8, 7, 9 from left to right). (**c**) Schematic of transplantation design, where each component can include middle-aged (M; 12mo) or young (Y; 2mo) sources. (**d**) Frequency of donor-derived (CD45.2+) myeloid (CD11b+) cells, B cells (B220+) and T cells (CD3+) in the blood at 24wks post-transplant (*n* = 7, 10, 6, 7 from left to right). (**b,d**) Dots represent individual mice and bars are mean ± SEM. All *n* values refer to the number of mice used. *P*-values were generated for (**b,d**) by one-way ANOVA with Holm-Sidak’s multiple comparisons test.

To further evaluate how myeloid-biased hematopoiesis in the peripheral blood related to alterations in the BM, we examined proportions of hematopoietic stem and progenitor cells (HSPCs) at 24 weeks post-transplant. It has been described that old HSCs transplanted into old recipient mice have reduced homing and ability to reconstitute the HSPC compartment and hematopoiesis^22^. Examining our middle-aged (MMM) versus young (YYY) control groups, we found that middle-aged LT-HSCs transplanted into middle-aged recipient mice have a reduced ability to reconstitute HSPC populations including LT-HSC, ST-HSC and MPP3/4 (Fig. 3A, C). Furthermore, examining the ratio of myeloid-restricted progenitors to LT-HSCs revealed greater proportion of myeloid progenitors relative to LT-HSCs in MMM versus YYY control groups (Fig. 3B, D), which correlates to myeloid-biased hematopoiesis observed in the peripheral blood. In our experimental group of young LT-HSCs transplanted into middle-aged recipient mice (YMM), we observed significantly reduced ST-HSC frequency (Fig. 3A) and increased ratio of myeloid-restricted progenitor cells to LT-HSCs (Fig. 3B) relative to YYY controls. In the reciprocal experimental group of middle-aged LT-HSCs transplanted into young recipient mice (MMY), we observed significantly increased ST-HSC frequency (Fig. 3C) and decreased ratio of myeloid-restricted progenitor cells to LT-HSCs (Fig. 3D). This led us to hypothesize that the young BM microenvironment may restore lineage-balanced differentiation of middle-aged LT-HSCs by reducing their myeloid-biased transcriptional program. To test this hypothesis, we performed RNA sequencing on donor-derived LT-HSCs purified from these transplants (Fig. 3E). We found that middle-aged LT-HSCs re-isolated from young recipients transcriptionally cluster with young controls using multidimensional scaling (Extended Data Fig. 3A). Of the 293 significantly differentially expressed genes (FDR < 0.05, FC > 1.5) in young versus middle-aged LT-HSCs re-isolated from control transplants, 102 of these were restored in expression following transplant of middle-aged LT-HSCs into the young recipient BM microenvironment (*P* < 1×10^−16^ using Fisher’s Exact Test) (Fig. 3E, Supplementary Table 2). Within these 102 genes, the most abundantly represented group downregulated in middle-aged LT-HSCs in a young microenvironment (MMY) were regulators of myeloid differentiation (ex. *Cybb, Ltf, Mpeg1*). Genes upregulated in middle-aged LT-HSCs in a young environment were involved in cellular metabolic processes (ex. *Pcbd1, Gpt2, Sult2b1*). At a pathway level, signatures that correlated to functional rejuvenation included reduction in myeloid differentiation, Cebpα network, and immune system response (Extended Data Fig. 3B). Taken together, these results suggest that at middle age, the BM microenvironment induces and is indispensable for myeloid-biased hematopoiesis.

**Fig. 3:**
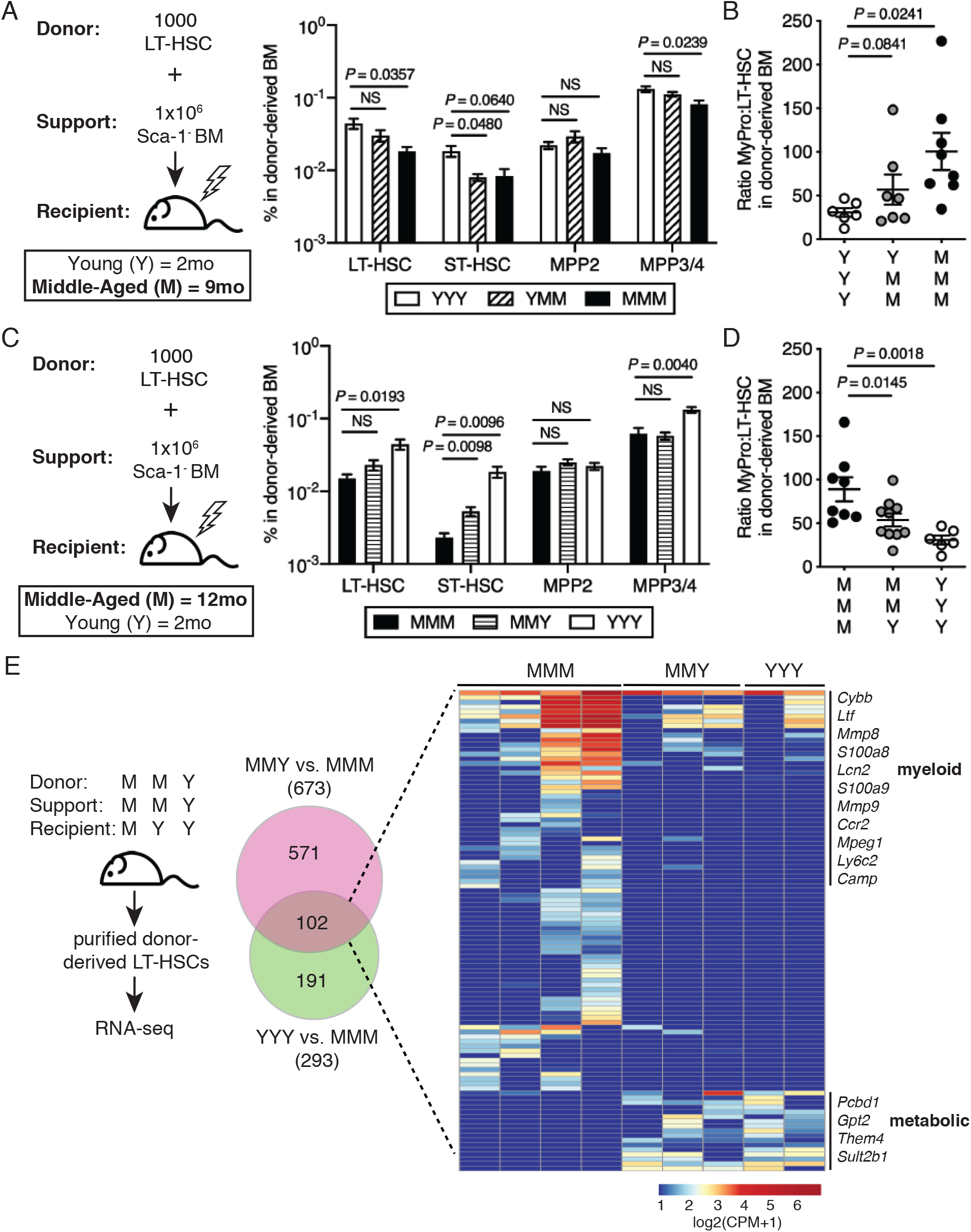
The young BM microenvironment is sufficient to reduce myeloid bias in middle-aged HSCs and progenitor cells. (**a**) Left: Schematic of transplantation design using purified LT-HSCs (CD45.2+) plus support cells (CD45.1+ Sca-1-) into recipient mice (CD45.1+), where each component can include young (Y; 2mo) or middle-aged (M; 9mo) sources. Right: Frequency of donor-derived LT-HSCs, ST-HSCs, MPP2 and MPP3/4 cells in the BM at 24wks post-transplant (*n* = 6, 7, 8, from left to right). (**b**) Ratio of donor-derived myeloid progenitors (MyPro; CD45.2+ cKit+ Sca-1-) to LT-HSCs (CD45.2+ SLAM) in the BM at 24wks post-transplant (*n* = 6, 7, 8, from left to right). (**c**) Left: Schematic of transplantation design, where each component can include middle-aged (M; 12mo) or young (Y; 2mo) sources. Right: Frequency of donor-derived LT-HSCs, ST-HSCs, MPP2 and MPP3/4 cells in the BM at 24wks post-transplant (*n* = 8, 10, 6, from left to right). (**d**) Ratio of donor-derived myeloid progenitors (MyPro) to LT-HSCs in the BM at 24wk post-transplant (*n* = 8, 10, 6, from left to right). (**e**) Left: Schematic of design to re-isolate donor-derived LT-HSCs for RNA-seq from the experiment shown in (c). Middle: Venn diagram of overlapping differentially expressed genes (*FDR* < 0.05, FC > 1.5) in middle-aged LT-HSCs in young recipients vs. middle-aged recipients, and young versus middle-aged LT-HSC transplant control groups. Right: Heatmap of expression of 102 overlapping genes in MMM, MMY and YYY conditions (*n* = 4, 3, 2, from left to right). (**a-d**) Dots represent individual mice and bars are mean ± SEM. All *n* values refer to the number of mice used. *P*-values were generated for (**a** and **c**) by Brown-Forsythe and Welch ANOVA with Dunnett’s T3 multiple comparisons test and for (**b** and **d**) by one-way ANOVA with Holm-Sidak’s multiple comparisons test.

Next, we used an unbiased approach to generate a list of candidate cytokines or growth factors that may be altered in the milieu of the middle-aged BM microenvironment. Our strategy at that time was to utilize known transcriptional changes between young and middle-aged multipotent progenitor cells from our published single-cell RNA-seq data^23^ as input into Ingenuity Pathway Analysis (IPA) Upstream Regulator analysis. This analysis predicts upstream cytokines and growth factors, increased or decreased, which would result in the observed transcriptional changes. This analysis predicted several upstream signaling molecules potentially altered in the middle-aged BM microenvironment including increased IGF2, and reduced NRG1, IGF1, TGFβ1 and EGF (Fig. 4A). We confirmed by IPA analysis of our RNA-seq data comparing young and middle-aged LT-HSCs (Fig. 1E, F) that IGF1, NRG1, IGF1, TGFβ1 and EGF were also predicted to be significantly enriched as upstream regulators (Supplementary Table 3). Reduced IGF1 in the middle-aged BM microenvironment became our top candidate to functionally test based on several experimental observations: (1) of these candidates, only *Igf1* and *Igf2* were found to be specifically expressed by non-hematopoietic cells in the BM microenvironment based on single-cell RNA-seq^24^ (Fig. 4B, Extended Data Fig. 4A), (2) decreased concentration of IGF1, but no change in IGF2, was observed in the BM fluid of middle-aged versus young mice (Fig. 4C, Extended Data Fig. 4B), (3) the highest expression of *Igf1* in the BM was observed in mesenchymal cells (Fig. 4B) and a bulk population of mesenchymal stromal cells (MSCs; CD45- Ter119- CD31- CD51+) (Fig. 4D), subsets of which have been previously identified as niches for hematopoietic stem and progenitor cells^25–28^, (4) *Igf1* expression in bulk MSCs was decreased in middle-aged compared to young MSCs (Fig. 4D), and (5) the major receptor for IGF1 signaling, *Igf1r*, was expressed in hematopoietic stem and progenitor cells (Extended Data Fig. 4C). We then experimentally evaluated whether genetic loss of IGF1 would be sufficient to cause hematopoietic aging phenotypes. We conditionally deleted *Igf1* in the environment by transplanting wild-type BM hematopoietic cells into tamoxifen-inducible *Igf1*-floxed mice (*Igf1*^fl/fl^; Cre-ER^T2^) and administering tamoxifen after recovery from transplantation to avoid the confounding role of *Igf1* in irradiation response^29^. We found that the *Igf1*-deficient environment increased myeloid cell production and decreased B lymphoid cell production from wild-type hematopoietic cells without altering overall engraftment level (Fig. 4E, Extended Data Fig. 4D). Conditional deletion of *Igf1r* on donor hematopoietic cells (*Igf1r*^fl/fl^; Mx1-Cre) transplanted into wild-type recipient mice also resulted in increased myeloid cell production and decreased B lymphoid cell production without altering overall engraftment level (Extended Data Fig. 4E). These data support a model that a decrease in IGF1 signaling, caused by either reduction in environment-produced IGF1 or reduction in expression of IGF1R on hematopoietic cells, is sufficient to cause myeloid-biased hematopoiesis.

**Fig. 4:**
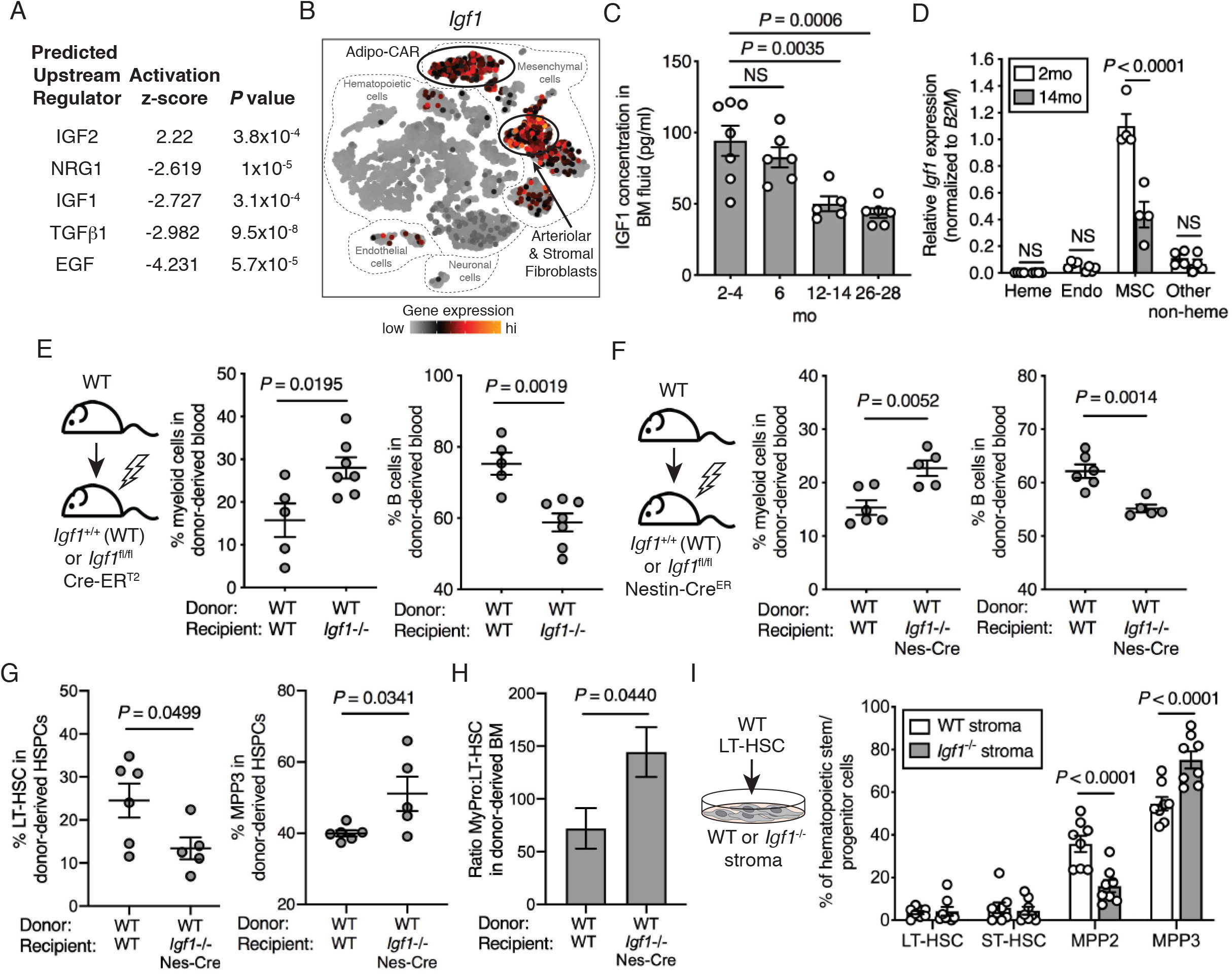
Reduction in local IGF1 from *Nestin*-expressing MSCs in the BM microenvironment causes myeloid-biased hematopoiesis. (**a**) Upstream growth factors predicted to be enriched in the middle-aged BM microenvironment by Ingenuity Pathway Analysis of middle-aged vs. young multipotent progenitor cell scRNA-seq data^23^. (**b**) Expression of *Igf1* in BM subsets assessed by scRNA-seq^24^. For detailed cell type annotation refer to: https://nicheview.shiny.embl.de. (**c**) IGF1 concentration in BM fluid of a single mouse femur from 2-4mo (*n* = 7), 6mo (n = 6), 12-14mo (*n* = 5) and 26-28mo (*n* = 6) mice. (**d**) Relative *Igf1* expression in hematopoietic (Heme; CD45+) cells, endothelial (Endo; CD45- Ter119- CD31+ CD51-) cells, MSCs (CD45- Ter119- CD31- CD51+) cells, and other non-heme (CD45- Ter119- CD31- CD51-) cells isolated from 2mo (*n* = 4) and 14mo (*n* = 4) mice. (**e**) Left: Experimental design. Right: Frequency of donor-derived myeloid cells (CD45.1+ CD11b+) and B cells (CD45.1+ B220+) in the blood at 24wks post-transplant of WT BM into lethally irradiated WT or *Igf1*^fl/fl^; CreER^T2^ recipient mice (*n* = 5, 7, from left to right). (**f**) Left: Experimental design. Right: Frequency of donor-derived myeloid cells (CD45.1+ CD11b+) and B cells (CD45.1+ B220+) in the blood at 20wks post-transplant of WT BM into lethally irradiated WT or *Igf1*^fl/fl^; Nestin-Cre^ER^ recipient mice (*n* = 6, 5, from left to right). (**g**) Frequency of donor-derived LT-HSCs and MPP3 cells in the BM at 20wks post-transplant of WT BM into lethally irradiated WT or *Igf1*^fl/fl^; Nestin-Cre^ER^ recipient mice (*n* = 6, 5, from left to right). (**h**) Ratio of donor-derived myeloid progenitors (Myel Pro; CD45.1+ cKit+ Sca-1-) to LT-HSCs (CD45.1+ SLAM) in the BM at 20wks post-transplant of WT BM into lethally irradiated WT or *Igf1*^fl/fl^; Nestin-Cre^ER^ recipient mice (*n* = 6, 5, from left to right). (**i**) Left: Experimental design. Right: Frequency of hematopoietic stem and progenitor populations after 4d co-culture of WT LT-HSCs with WT or *Igf1*^−/-^ stroma (*n* = 3, done in replicate or triplicate 3 independent times). (**c**-**h)** Dots represent individual mice and bars are mean ± SEM. (**i**) Dots represent replicates from pooled experiments repeated in triplicate. *P*-values were generated for (**c**) by one-way ANOVA with Holm-Sidak’s multiple comparisons test, (**d**, **i**) by two-way ANOVA with Sidak’s multiple comparisons test, (**e**-**g**) by unpaired, two-tailed *t* tests.

To discern the effects of systemic (liver-derived) versus local (BM microenvironment-derived) IGF1 on hematopoiesis, we utilized an inducible *Nestin*-Cre^ER^ model^30,31^. For this experiment, an inducible Cre model was critical given the embryonic lethality of *Igf1* knockout mice^32^ and the importance of *Igf1* in skeletal patterning during development^33^, which could confound interpretation of results when using Cre drivers in the BM microenvironment that are expressed developmentally. We transplanted wild-type BM hematopoietic cells into *Igf1*^fl/fl^; *Nestin*-Cre^ER^ mice and administered tamoxifen after recovery from transplantation. We found that transplant into *Igf1*^fl/fl^; *Nestin*-Cre^ER^ recipients increased myeloid cell production and decreased B lymphoid cell production without altering overall engraftment level (Fig. 4F, Extended Data Fig. 4F). As the liver is the main source of circulating IGF1^35^ and *Igf1*^fl/fl^; *Nestin*-Cre^ER^ mice do not induce deletion of *Igf1* in the liver (Extended Data Fig. 4G), we conclude that decrease in IGF1 in the local BM microenvironment is sufficient to cause myeloid-biased hematopoiesis.

To further evaluate the direct role of BM microenvironment-derived IGF1 on hematopoietic stem and progenitor cells, we analyzed the BM of our transplanted *Igf1*^fl/fl^; *Nestin*-Cre^ER^ mice at 24 weeks post-transplant. Similar to the phenotype observed upon transplantation of young LT-HSCs into middle-aged recipient mice (Fig. 3A, B), we observed that transplant into *Igf1*^fl/fl^; *Nestin*-Cre^ER^ mice resulted in decreased frequency of LT-HSCs (Fig. 4G), increased frequency of the myeloid-biased multipotent progenitor MPP3, and increased the ratio of myeloid-restricted progenitor cells to LT-HSCs (Fig. 4H). Lastly, we performed short-term co-culture of wild-type, purified LT-HSCs seeded onto *Igf1*^fl/fl^; Cre-ER^T2^ BM-derived stroma (Fig. 4I). We found that co-culture with *Igf1*-deficient BM-derived stroma caused enhanced production of phenotypic myeloid-biased multipotent progenitor (MPP3) cells at the expense of erythroid-biased MPP2 cells^36^, assessed by cell surface marker staining and flow cytometry. Together, these results support that decreased production of IGF1 by the middle-aged BM microenvironment is sufficient to cause myeloid-biased hematopoiesis and directly impacts HSPCs.

We next asked whether stimulation of middle-aged LT-HSCs with IGF1 could rejuvenate these cells at the functional and molecular levels. We isolated and treated middle-aged LT-HSCs with recombinant IGF1 *in vitro* for 20 min and found increased phosphorylation of IGF1R and downstream AKT (Fig. 5A,B), supporting that these cells are capable of directly responding to IGF1. IGF1 stimulation of middle-aged LT-HSCs for two days *in vitro* resulted in reduced myeloid differentiation (Extended Data Fig. 5A). We next tested the effect of IGF1 directly on middle-aged LT-HSCs using a polyvinyl alcohol (PVA)-based culturing method recently shown to maintain and expand functional HSCs^37^, followed by transplantation into recipient mice (Fig. 5C). A single spike-in of recombinant IGF1 at the start of these cultures resulted in decreased myeloid cell differentiation and increased B lymphocyte differentiation *in vivo*, without altering overall engraftment level (Fig. 5C, Extended Data Fig. 5B). At a molecular level, short-term (18h) *in vitro* IGF1 stimulation decreased the frequency of middle-aged LT-HSCs with γH2AX foci (Fig. 5D), and increased polarity of CDC42 and tubulin (Fig. 5E). To examine the immediate transcriptional target genes activated or repressed by IGF1 signaling in middle-aged LT-HSCs, we performed RNA-seq after an 18h *in vitro* stimulation with recombinant IGF1 (Fig. 5F). IGF1 stimulation resulted in differential expression of 154 genes (*P (unadj)* < 0.015, FC > 1.5) in middle-aged LT-HSCs. Of these, 32 genes were found to be differentially expressed in middle-aged versus young, vehicle-treated LT-HSCs (*P* < 1×10^−16^ using Fisher’s Exact Test), including *Sla, CD74, Spi1* (PU.1), and *Fancc* (Supplementary Table 4). At a pathway level, IGF1 stimulation of middle-aged LT-HSCs rescued target genes enriched for PI3K/AKT/mTOR signaling, G2M checkpoint, and xenobiotic metabolism (Fig. 5G), and increased expression of genes enriched in epigenetic-related signatures (Extended Data Fig. 5C). In addition, IGF1 stimulation decreased myeloid-biased LT-HSC and increased lymphoid-biased LT-HSC signatures (Extended Data Fig. 5D). Of note, several pathways were not found to be rescued upon IGF1 stimulation including increased TNFα signaling and IFNγ response, uncoupling these inflammatory pathways from the observed rejuvenation of middle-aged LT-HSCs. Taken together, these results suggest that enhanced local IGF1 signaling at middle age can rejuvenate molecular and functional hallmarks of LT-HSC aging.

**Fig. 5:**
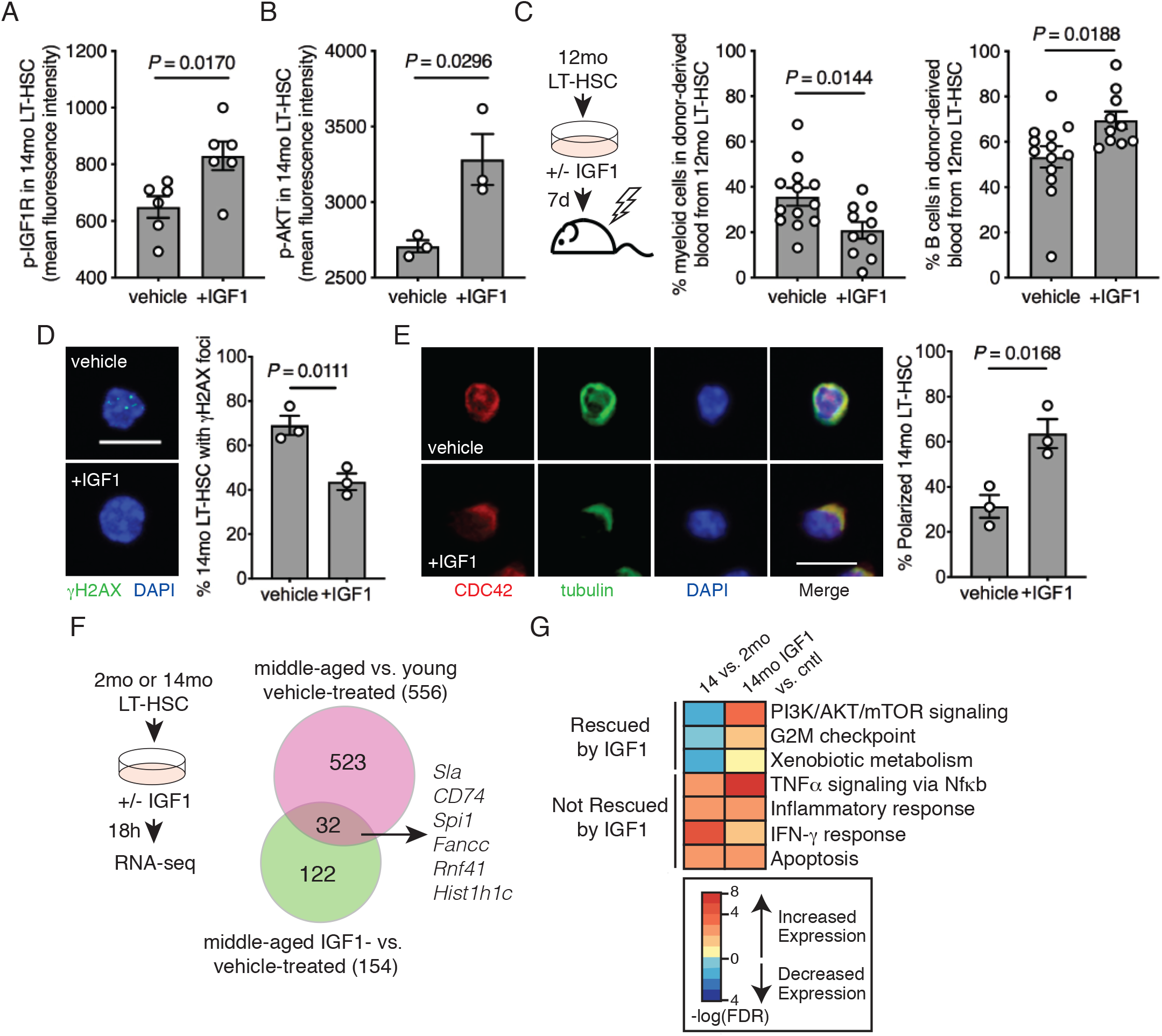
IGF1 stimulation of middle-aged LT-HSCs rescues hallmarks of aging. (**a, b**) Quantification of intracellular phospho-flow cytometry analysis of (**a**) p-IGF1R (*n* = 6) and (**b**) p-AKT (*n* = 3) following stimulation of 14mo LT-HSCs with IGF1 or vehicle. (**c**) Left: Experimental design for 7d *ex vivo* stimulation of 12mo LT-HSC with IGF1 or vehicle followed by transplant. Center: Frequency of donor-derived myeloid cells (CD45.2+ CD11b+) in the blood of recipient mice (*n* = 13, 10 left to right). Right: Frequency of donor-derived B cells (CD45.2+ B220+) in the blood of recipient mice at 8wks post-transplant (*n* = 13, 10 left to right). (**d**) Left: Representative images of γH2AX and DAPI-stained 13-14mo LT-HSCs stimulated with IGF1 or vehicle. Scale bar, 10 μm. Right: Quantification of the percentage of 13-14mo LT-HSC with γH2AX foci after stimulation with vehicle (*n =* 3) or IGF1 (*n =* 3). (**e**) Left: Representative images of 13-14mo LT-HSCs stimulated with vehicle or IGF1 and stained with CDC42, tubulin, DAPI, and overlay. Scale bar, 10 μm. Right: Quantification of the percentage of 13-14mo LT-HSC with polarized CDC42 and tubulin after stimulation with vehicle (*n =* 3) or IGF1 (*n =* 3). (**f**) Left: Experimental design for 18hr *in vitro* stimulation of 2mo and 14mo LT-HSC with IGF1 or vehicle followed by RNA-seq. Right: Venn diagram of overlapping differentially expressed genes in 14mo vs. 2mo LT-HSC and 14mo LT-HSC stimulated with IGF1 vs. vehicle. (**g**) Heatmap of gene set enrichment analysis of 14mo vs. 2mo LT-HSCs, and 14mo LT-HSCs stimulated with IGF1 vs. vehicle. (**a**-**e**) Dots represent individual mice and bars are mean ± SEM. All *n* values refer to the number of mice used. *P*-values were generated for (**a**-**e**) by unpaired two-tailed *t*-test.

## Discussion

Here, we have discovered that processes causing impaired hematopoietic function with aging have already been initiated by middle age, including changes in the non-hematopoietic BM microenvironment. This not only establishes the BM microenvironment as a critical component causing hematopoietic aging but also suggests that targeting the BM microenvironment at middle age has strong potential to rejuvenate HSCs and hematopoiesis. Previous studies performing binary comparisons between young and old individuals^1,2,4–8^ have led to conflicting results with respect to whether HSC-intrinsic or -extrinsic alterations drive aging. Our work supports that, at the time in which some of the hallmarks of aging are first observed, HSC-extrinsic alterations are indispensable for and induce these phenotypes.

Previous work by several groups has demonstrated that old HSC function is not improved by transplantation into young recipient mice^38–40^, suggesting that there is an upper limit on the age at which rejuvenation of aging HSCs by a young BM microenvironment is possible. Our data, both at a functional and transcriptional level, define middle-aged LT-HSCs as being in a unique state where they exhibit some of the hallmarks of HSC aging but remain responsive to extrinsic effects from the BM microenvironment. So, what defines this ‘window of opportunity’ for rejuvenation? While a drop in local IGF1 levels in the milieu of the BM may be potentially used to define the lower boundary, we have also observed that IGF1 levels do not decrease further in old mice and thus cannot be utilized to define the upper boundary of rejuvenation potential, in addition to this being an impractical measurement in live organisms. The transcriptome profiling datasets that we have generated in this study comparing young, middle-aged and old LT-HSCs are an important foundation to begin to predict and evaluate surrogate markers of HSC rejuvenation potential that may be more readily assessed in live organisms. At a mechanistic level, identification of epigenetic-related signatures induced by IGF1 stimulation of middle-aged LT-HSCs supports an intriguing hypothesis that the ‘window of opportunity’ for rejuvenation may be related to the level of HSC-intrinsic epigenetic plasticity.

At a mechanistic level, we have discovered that myeloid-biased differentiation, one of the hallmarks of LT-HSC aging, is initiated by declining levels of IGF1 in the local BM microenvironment. While we found reduced myeloid-biased differentiation by stimulating with recombinant IGF1 *ex vivo*, IGF1 alone did not fully recapitulate the restored HSC function that was achieved by transplanting middle-aged LT-HSCs into a young BM microenvironment. This suggests that additional factors may play a key role in this process. Alternatively, recombinant IGF1 may be insufficient in effectively activating the same downstream pathway(s) that are achieved by locally produced IGF1 *in vivo*. In muscle, heart and skin, it has been demonstrated that local IGF1 plays an autocrine/paracrine role in promoting regeneration and is derived from distinct IGF1 pre-propeptides, distinct from those which contribute to liver-derived, circulating IGF1 levels^41^. In contrast to our reported findings, previous work has demonstrated that both fasting and dietary restriction reduce circulating IGF1 levels and result in beneficial effects on HSC function^42,43^. This suggests that variables including systemic versus local IGF1, age, and dietary status are relevant variables to be considered in the design of potential prophylactic therapies. We suggest that developing approaches to therapeutically target MSCs to restore local production of IGF1 in the BM microenvironment, or methods to increase downstream AKT or mTOR signaling in middle-aged HSCs, are compelling strategies to preserve healthy function of the hematopoietic system from middle age into older age.

Although our study focuses on and provides novel insights into changes that occur in hematopoietic cells at middle age, the changes occurring in the middle-aged BM microenvironment that contribute to these phenotypes have not yet been comprehensively assessed. In particular, while we have defined that *Igf1* expression is reduced in a bulk, heterogeneous population of MSCs, and that loss of *Igf1* in *Nestin*-expressing cells in the BM microenvironment can recapitulate middle-aged hematopoiesis phenotypes, we have not comprehensively determined whether there are other subsets of non-*Nestin*-expressing cells that produce *Igf1* and/or play a role in aging HSPC function. Future studies to examine MSC heterogeneity and changes in expression of soluble factors by single-cell RNA-seq, spatial transcriptomics, and lineage tracing will be the ideal approaches to address these questions.

Much effort and emphasis has been placed on understanding the role of inflammation in driving aging phenotypes (“inflamm-aging”). The transcriptional studies presented here of young, middle-aged and old LT-HSCs suggest that the impact of inflammation on LT-HSCs is likely progressive and cumulative with age. Our study also demonstrates that rejuvenation of some of the hallmarks of LT-HSC aging at middle age can occur without altering inflammation signatures. This suggests that it may be possible to tease apart pro-regenerative and HSC lineage commitment pathways from inflammation. Whether promoting IGF1 signaling while simultaneously reducing inflammation would have further benefit in extending hematopoietic healthspan remains to be tested.

## Methods

### Mice

C57BL/6J female and male mice and B6.SJL-*Ptprc^a^Pepc^b^*/BoyJ (referred to as B6.CD45.1) mice were obtained from, and aged within, The Jackson Laboratory. Young mice ranged from 2-4mo and middle-aged mice ranged from 9-14mo for experiments, all other ages used are noted in the figures. B6.129(FVB)-*Igf1^tm1Dlr^*/J (referred to as Igf1^fl/fl^) were crossed to B6.129-*Gt(ROSA)26Sor^tm1(cre/ERT2)Tyj^*/J (referred to as Cre-ER^T2^) and were crossed to C57BL/6-Tg(Nes-cre/Esr1*)1Kuan/J (referred to as Nestin-Cre^ER^, and B6;129-*Igf1r^tm2Arge^*/J (referred to as Igf1R^fl/fl^) were crossed to B6.Cg-Tg (Mx1-cre)1Cgn/J mice (referred to as Mx1-Cre). To induce Mx1-Cre recombinase expression, mice were injected once, every other day for five days, by intraperitoneal (IP) injections of 15□mg/kg high molecular weight polyinosinic-polycytidylic acid (pIpC) (InvivoGen). To induce Cre-ER^T2^ recombinase expression mice received 125□mg/kg tamoxifen for three consecutive days by oral gavage. For liver samples and control, after tamoxifen administration, genomic DNA was extracted from liver, and BM cells for control, for recombination PCR using specified primers (Supplementary Table 5). All mouse experiments and protocols were approved by The Animal Care and Use Committee at The Jackson Laboratory.

### Peripheral blood analysis

Blood was collected from mice via retro-orbital sinus and red blood cells were lysed before staining with CD45.1 (clone A20), CD45.2 (clone 104), B220 (clone RA3-6B2), CD3e (clone 145-2C11), CD11b (clone M1/70), Ly6g (clone 1A8), Ly6c (clone HK1.4), GR-1 (clone RB6-8C5). Stained cells were analyzed on an LSRII (BD) and populations were analyzed using FlowJo V10. CBCs were performed on collected blood using an Advia 120 Hematology Analyzer (Siemens).

### Isolation and phenotyping of hematopoietic stem and progenitor cells

BM mononuclear cells (MNCs) were prepared from pooled and crushed femurs, tibiae, and iliac crests of each individual mouse. MNCs were isolated by Ficoll-Paque (GE Healthcare Life Sciences) density centrifugation and stained with a combination of fluorochrome-conjugated antibodies: c-Kit (clone 2B8), Sca-1 (clone D7), CD150 (clone TC15-12F12.2), CD48 (clone HM48-1), CD34 (clone RAM34), CD41 (clone MWReg30), EPCR (clone RMEPCR1560), FLT3 (clone A2F10), mature lineage (Lin) marker mix (B220 (clone RA3-6B2), CD11b (clone M1/70), CD4 (clone RM4-5), CD5 (clone 53-7.3), CD8a (clone 53-6.7), Ter-119 (clone TER-119), Gr-1 (clone RB6-8C5)), and the viability stain propidium iodide (PI). For LT-HSC phenotyping, cell frequency was determined based on the following surface marker profiles using a FACSymphony A5 (BD): LT-HSC (SLAM) (Lin- Sca-1+ c-Kit+ Flt3- CD150+ CD48-), LT-HSC (EPCR) (Lin- Sca-1+ c-Kit+ CD34- EPCR+), LT-HSC (CD41+) (Lin- Sca-1+ c-Kit+ Flt3- CD150+ CD48- CD41+). LT-HSCs were isolated for all experiments using a FACSAria II (BD) based on SLAM markers. All flow cytometry data was analyzed using FlowJo V10 software.

### Immunofluorescence Staining of LT-HSCs

Staining for γH2AX or CDC42 and tubulin was performed as previously described^13,44^. As defined by SLAM markers, 2,000 LT-HSCs were sorted directly into 96 well plates containing SFEMII with Pen-Strep and SCF (100 ng/ml), TPO (50 ng/ul), with and without IGF1 (100 ng/ml), (BioLegend, Peprotech and StemCell Technologies) for 16-18 hrs at 37°C and 5% CO_2_. LT-HSCs were plated directly onto retronectin-coated coverslips and allowed to adhere for 2 hrs at 37°C and 5% CO_2_. Cells were fixed in 4% PFA (Affymetrix) for 15 mins at 4°C, and permeabilized with 0.2% Triton X-100 (Fisher Scientific) in PBS for 20 mins at room temperature (RT) and blocked with 10% goat serum (Life Technologies) for 20 mins at RT. LT-HSCs were stained with conjugated γH2AX antibody (FITC-conjugated anti-phospho-H2A.X antibody, Biolegend, clone 2F3, 1:50 dilution) for 2 hrs at 37°C, washed 3x with PBS and mounted on slides with Gold Antifade with DAPI (Invitrogen). LT-HSCs were stained with rabbit polyclonal anti-CDC42 antibody (Millipore, 1:50 dilution) and rat monoclonal anti-tubulin antibody (Abcam, 1:100 dilution) for 2 hrs at 37°C, then washed 3x with PBS before secondary antibodies, α-Rabbit conjugated with Alexa-568 (Invitrogen, 1:1000 dilution) and α-rat conjugated with Dylight488 (Jackson Immuno Research Inc., 1:100 dilution) for 1 hr. Coverslips were mounted on slides with Gold Antifade with DAPI (Invitrogen). Images were acquired using a Leica SP8 confocal microscope and z-stack images were analyzed using Fiji software. Scale bars in images represent 10μm.

### RNA-seq library prep and analysis

For RNA-seq experiments shown in Fig. 1, young (2mo) and middle-aged (12mo) and old (22mo) LT-HSCs from 3 independent replicates of pooled mice were sorted directly into RLT buffer. For RNA-seq experiments in Fig. 3, donor-derived (CD45.2+) LT-HSCs were re-isolated from 2-4 independent replicates of transplanted mice (YYY, MMY and MMM) and sorted directly into RLT buffer. For RNA-seq experiments shown in Fig. 5, young (2mo) and middle-aged (14mo) LT-HSCs from 3 independent replicates of pooled mice were sorted directly into a 96-well plate for 18hrs in StemSpan SFEM II, SCF (100 ng/ml), TPO (50 ng/ml), Pen-Strep (1%) with vehicle (0.1% BSA in PBS) or IGF1 (100ng/ml) stimulation at 37°C and 5% CO_2_ before being centrifuged and resuspended in RLT buffer. Total RNA was isolated from cells using the RNeasy Micro kit (Qiagen). Sample quality was assessed using the Nanodrop 2000 spectrophotometer (Thermo Scientific) and the RNA 6000 Pico LabChip assay (Agilent Technologies). Libraries were prepared by the Genome Technologies core facility at The Jackson Laboratory using the Ovation RNA-seq System V2 (NuGEN Technologies) and Hyper Prep Kit (Kapa Biosystems). Libraries were checked for quality and concentration using the D5000 ScreenTape assay (Agilent Technologies) and quantitative PCR (Kapa Biosystems), according to the manufacturers’ instructions. Libraries for RNA-seq experiments shown in Fig. 1 and Fig. 4, were pooled and sequenced 75 bp paired-end on the NextSeq 500 (Illumina) using NextSeq High Output Kit v2.5 reagents (Illumina). Libraries for RNA-seq experiments shown in Fig. 2, were pooled and sequenced 75 bp single-end on the NextSeq 500 (Illumina) using NextSeq High Output Kit v2 reagents (Illumina). Raw and processed data are publicly available in the Gene Expression Omnibus (GSE144933, GSE144934 and GSE151333). Trimmed alignment files (with trimmed base quality value < 30, and 70% of read bases surpassing that threshold) were processed using the RSEM (v1.2.12; RNA-Seq by Expectation-Maximization) software and the Mus Musculus reference GRCm38. Alignment was completed using Bowtie 2 (v2.2.0) and processed using SAMtools (v0.1.18). Expected read counts per gene produced by RSEM were rounded to integer values, filtered to include only genes that have at least two samples within a sample group having counts per million reads (cpm) > 1.0, and were passed to edgeR (v3.5.3) for differential expression analysis. A negative binomial generalized log-linear model was fit to the read counts for each gene.

The dispersion trend was estimated by Cox-Reid approximate profile likelihood followed by empirical Bayes estimate of the negative binomial dispersion parameter for each tag, with expression levels specified by a log-linear model. Likelihood ratio tests for coefficient contrasts in the linear model were evaluated producing a p-value per contrast. The Benjamini and Hochberg’s algorithm was used to control the false discovery rate (FDR). Unless otherwise, indicated, features with a fold change (FC) > 1.5 and an FDR-adjusted *P*-value < 0.05 were declared significantly differentially expressed. Gene set enrichment analysis (GSEA)^45^ was performed using previously published human HSC and mouse LT-HSC RNA-seq data^2,15^. GSEA Molecular Signatures Database (MSigDB) was utilized to analyze differential expression of hallmark gene sets, curated gene sets and gene ontology gene sets (http://software.broadinstitute.org/gsea/msigdb/index.jsp). Clusters of genes in Fig. 1 were derived from the input of all differentially expressed genes (DEGs) in the denoted comparisons and compiled based on similarity of gene expression patterns across the three conditions. In detail, for each gene, mean cpm of young, middle-aged and old were calculated. Fold change for each gene was calculated from young to middle-aged, and middle-aged to old by dividing the mean cpm values. A cutoff fold change of 1.2 was used to classify groups of genes that were altered in expression (FC > 1.2), or not altered in expression (FC < 1.2), from young to middle-aged, and/or middle-aged to old. Ingenuity Pathway Analysis (Ingenuity Systems; https://www.qiagenbioinformatics.com/products/ingenuity-path-way-analysis/) is a computational tool that uses prior knowledge of expected effects between cytokines or growth factors and their target genes. This analysis was used to examine how many known target genes of each cytokine or growth factor were present in our young versus middle-aged multipotent progenitor single-cell RNA-seq data^24^, and also compared the direction of change to what was expected from the literature, to predict likely relevant factors. *P-*values measuring statistical significance were determined by the software using Fisher’s Exact Test.

### *In vivo* transplantation

For aging transplantation experiments, B6.CD45.1 female recipient mice (2mo, 9mo or 12mo) were lethally irradiated (12Gy gamma irradiation, split dose). 1,000 LT-HSCs from C57BL/6J mice (2mo, 9mo or 12mo) together with 1×10^6^ sorted Sca-1-BM support cells from B6.CD45.1 mice (2mo, 9mo or 12mo) were transplanted intravenously by retro-orbital injection. For the 9mo and 12mo dataset, 5 separate sorts were performed for each to isolate the required cell numbers for donor, support and recipient groups. Each group was comprised from multiple donors and received experimental mice from multiple sort days. For IGF1 mouse model transplantation experiments, 2×10^6^ MNCs from one donor B6.CD45.1 mouse were transplanted into indicated numbers of lethally irradiated *Igf1*^+/+^ Cre-ER^T2^ mice and *Igf1*^fl/fl^ Cre-ER^T2^ recipient mice on each of two separate transplant days, or from one donor B6.CD45.1 mouse into indicated numbers of *Igf1*^+/+^ Nestin-Cre^ER^ and *Igf1*^−/-^ Nestin-Cre^ER^ mice on one transplant day. For IGF1R mouse model transplantation experiments, LT-HSC (SLAM) from two *Igf1R*^fl/fl^ Mx1-cre donor mice or three C57BL/6J control donor mice were sorted into a 96-well plate and expanded for 48 hrs before harvesting the well and transplanting with 2×10^6^ (CD45.1+) MNCs support cells into lethally irradiated B6.CD45.1. *Igf1R*^fl/fl^ Mx1-cre received pIpC injections four weeks before transplant and Cre-ER^T2^ and Nestin-Cre^ER^ recipient mice received tamoxifen four weeks post-transplant, as described above. For IGF1 *ex vivo* treatment of LT-HSCs followed by transplantation experiments, 50 LT-HSC isolated from three pooled C57BL/6J donor mice, after seven days in culture, were mixed with 1×10^6^ MNCs from one B6.CD45.1 mouse into indicated numbers of lethally irradiated B6.CD45.1 recipient mice, each in two separate transplant days. Multilineage peripheral blood reconstitution was monitored every four weeks thereafter by flow cytometry analysis of blood samples using a cocktail of CD45.1, CD45.2, CD11b, B220, CD3∊, and Gr-1 on an LSRII (BD). CBCs were performed on peripheral blood samples at 24 weeks post-transplant using an Advia 120 Hematology Analyzer (Siemens).

### IGF1 expression patterns in the BM

The expression pattern of *Igf1, Igf2, Nrg1, Tgfb1, Egf and Ig1r* in hematopoietic stem and progenitor cells and BM microenvironment cells was analyzed utilizing single-cell RNA-seq data obtained from the BM of young WT C57BL/6 mice^24^.

### IGF1 and IGF2 concentrations in the BM

BM fluid was flushed and isolated from single femurs of female C57BL/6J mice from ages 2 mo to 28 mo. A mouse-specific IGF1 Immunoassay kit (ELISA, R&D Systems) and a mouse-specific IGF2 PicoKine kit (ELISA, Boster biological technology) was used to determine the concentration of IGF1 and IGF2 proteins in the BM.

### Real-Time PCR

Methods for isolation of MSCs, other BM microenvironment and hematopoietic cells were adapted from^46^. Briefly, BM plugs were flushed and digested in HBSS buffer containing collagenase type IV (2 mg/ml, GIBCO) and dispase (1mg/ml, Sigma) 3x for 10 mins at 37°C and 5% CO_2_. Each digested supernatant fraction was collected and added to ice-cold FACs buffer. Cell populations were then stained and sorted on a FACSAria II (BD) as follows: hematopoietic cells (Heme; CD45+), endothelial cells (Endo; CD45- Ter119- CD31+), mesenchymal stromal cells (MSC; CD45- Ter119- CD31- CD51+), and other BM microenvironment cells (other non-heme; CD45- Ter119- CD31- CD51-). RNA was isolated using RNeasy microkit or RNeasy minikit (Qiagen) and quantitative PCR was performed using RT^2^ Green ROX qPCR Mastermix (Qiagen) on Viaa7 (Applied Biosystems) or QuantStudio 7 Flex (Applied Biosystems). All mRNA expression levels were calculated relative to the housekeeping gene, *B2M*. *B2M* was chosen as our reference datasets showed no changes in this gene with age. Primer sequences for real-time PCR can be found in Supplementary Table 5.

### Colony-forming unit (CFU) assay

LT-HSCs were isolated and plated in liquid culture with vehicle (0.1%BSA/PBS) or with IGF1 (100ng/ml) for 48 hr before plating in MethoCult GF M3434 (StemCell Technologies) at the indicated numbers and cultured at 37°C and 5% CO_2_. Colonies were scored between 6- and 14-days post-plating using a Nikon Eclipse TS100 inverted microscope.

### Isolation and co-culture of MSCs

MSCs isolation and culture was adapted from^47^. Briefly, femurs, tibiae and iliac crest were isolated from 2 mo *Igf1^+/+^* Cre-ER^T2^ mice and *Igf1^fl/fl^* Cre-ER^T2^ mice and bones were flushed with α-MEM/15% FBS into 10 cm tissue culture treated dishes. Cells were allowed to adhere and grow at 37°C and 5% CO2 for five days. On day five, cells were split 1:3 and allowed to grow to confluence before being split again. MSCs were then used for experiments and not used beyond P5. For co-cultures, MSCs were seeded at 10,000 cells/well in a 48-well plate for 24 hrs to adhere. MSCs were treated with 20 μM 4-hydroxytamoxifen (4-OHT) for 24 hrs to activate Cre-ER^T2^ MSCs and washed 1× before adding 200 purified LT-HSC (Lin− Sca-1+ c-Kit+ Flt3− CD150+ CD48-) sorted into SFEMII, SCF (10ng//ml), IL3 (10ng/ml), IL6 (10ng/ml), and 1% Pen-Strep. Cells were collected and analyzed by flow cytometry on a FACSymphony A5 (BD) four days later.

### Phospho-flow cytometry

Middle-aged LT-HSCs from individual mice were sorted and resuspended in StemSpan SFEM II and divided for stimulation with vehicle (0.1% BSA in PBS) or IGF1 (100ng/ml) for 20 mins at 37°C. LT-HSCs were immediately fixed in 16% PFA for 10 mins at RT. Cells were then permeabilized in ice-cold 100% acetone for 10 mins on ice. Fixed and permeabilized cells were stained with intracellular phospho-IGF1R (pY1131) at 3:100 or phospho-AKT (pS473) at 1:20 dilutions for 30 mins at RT before analysis on FACSAria II (BD).

### *In vitro* LT-HSC culture with PVA and fibronectin

Methods for maintaining and transplanting LT-HSCs and stimulating with IGF1 were adapted from^37^. Briefly, 50 LT-HSCs (Lin- c-Kit+ Sca-1+ CD34- CD150+) were plated in a single well of a 96-well fibronectin-coated plate and cultured in 200 ul of medium composed of F12, 1% ITSX, 10 mM HEPES, 1% P/S/G, TPO (100 ng/ml), SCF (10 ng/ml) and PVA (1 mg/ml) with vehicle (0.1%BSA in PBS) or IGF1 (100ng/ml) at 37°C and 5% CO_2_ for seven days before transplant. For transplant, one well was harvested and combined with 1×10^6 BM competitor cells from B6.CD45.1 mice and transplanted into a single lethally-irradiated B6.CD45.1 recipient mouse. Peripheral blood was analyzed 8wks post-transplant.

## Acknowledgements

This work was supported by National Institute of Diabetes and Digestive and Kidney Diseases (NIDDK) grants R56DK112947 and R01DK118072, and the Ellison Medical Foundation New Scholar Award in Aging (J.J.T.). K.Y. received support from the Eunice Kennedy Shriver National Institute of Child Health and Human Development (NICHD) T32HD007065 and the American Society of Hematology (ASH) Scholar Award. Research reported in this publication was partially supported by the National Cancer Institute (NCI) grant P30CA034196 and National Institute of Aging (NIA) grant P30AG038070. We thank Nicole Dean, Jennifer SanMiguel, Kristina Mujica, Logan Schwartz and Amy LoTemplio for technical help, experimental and laboratory support, and Eric Pietras, Ross Levine, Julia Maxson, and Nadia Rosenthal for helpful discussion and critical comments. We thank Scientific Services at The Jackson Laboratory including: Flow Cytometry (Will Schott, Erin Kitten, Danielle Littlefield), Genome Technologies, and Microscopy.

## Author Contributions

K.Y. and J.J.T. designed experiments. K.Y., E.E., R.K.B., and M.A.L. performed and analyzed experiments. S.H. and L.V. provided analyzed data. K.Y. and J.J.T. wrote the manuscript. E.E. edited the manuscript.

---

**Extended Data Fig. 1:**
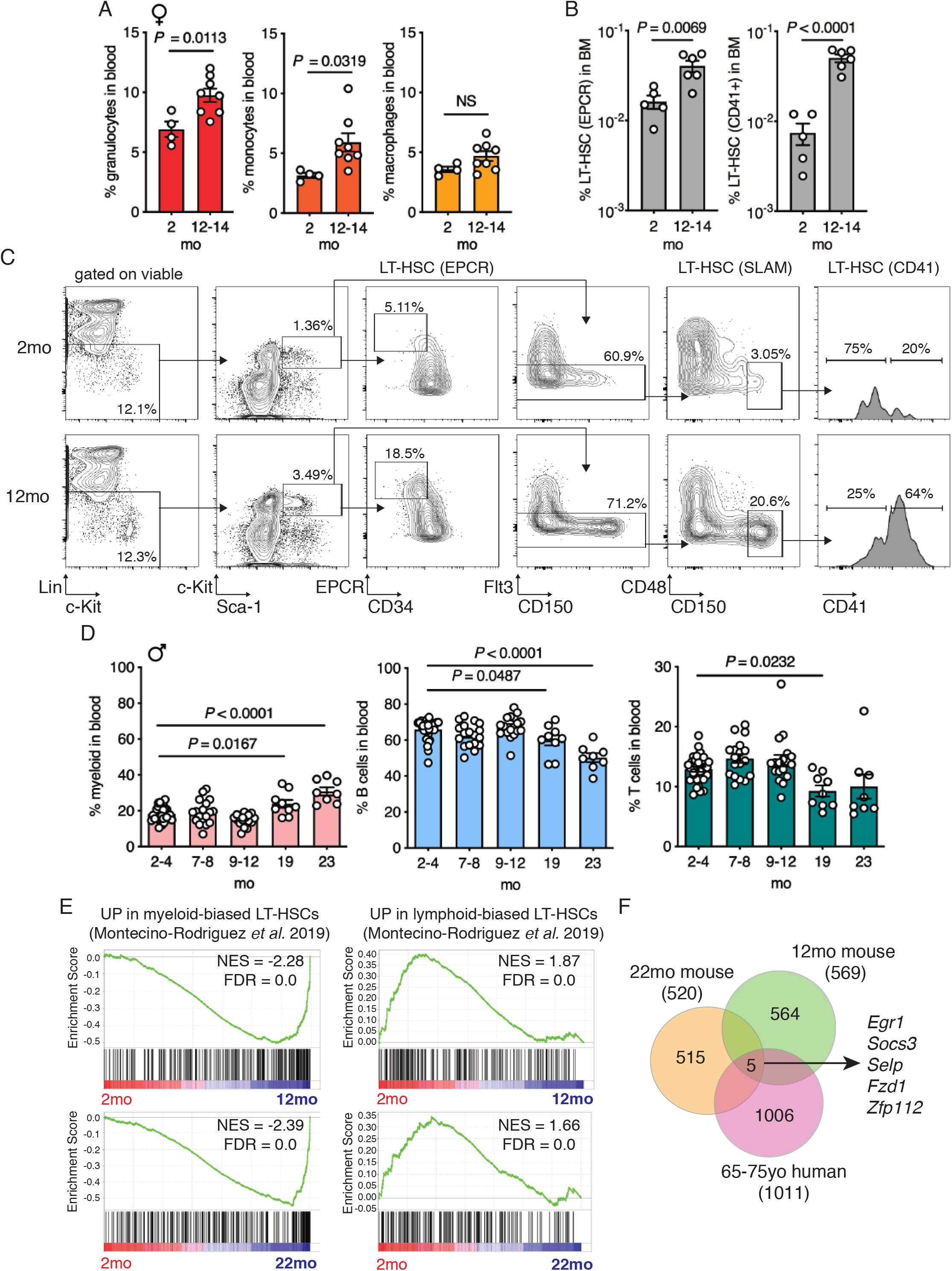
LT-HSC aging signatures are observed by middle age in mice. (**a**) Frequency of granulocytes (CD11b+ Ly6c+ Ly6g+), monocytes (CD11b+ Ly6c+ Ly6g-) and macrophages (CD11b+ Ly6c-Ly6g+) within the blood of female mice at 2mo (*n* = 4) and 12-14mo (*n* = 8). (**b**) Frequency of LT-HSCs defined by markers including EPCR (left) and CD41 (right) in whole BM of mice at 2mo (*n =* 5) and 12-14mo (*n =* 6). (**c**) Flow cytometry gating strategy showing frequency of LT-HSC populations in representative 2mo and 12mo female mice. (**d**) Frequency of myeloid cells (CD11b+), B cells (B220+) and T cells (CD3+) within the blood of male mice at 2-4mo (*n =* 30), 7-8mo (*n =* 9), 9-12mo (*n =* 18), 19mo (*n =* 9) and 23mo (*n =* 9). (**e**) Left: Enrichment of a myeloid-biased LT-HSC signature^15^ in 12mo vs. 2mo LT-HSCs (top) and in 22mo vs. 2mo LT-HSCs (bottom). Right: Enrichment of a lymphoid-biased LT-HSC signature^15^ in 2mo vs. 12mo LT-HSCs (top) and in 2mo vs. 22mo LT-HSCs (bottom). NES; normalized enrichment score. FDR; false discovery rate. (**f**) Venn diagram of overlapping differentially-expressed genes (*P* < 0.05, FDR > 1.5) in aged human HSCs (65-75yo), middle-aged mouse LT-HSCs (12mo) and old mouse LT-HSCs (22mo) compared to their respective young controls. (**a**, **b, d**) Dots represent individual mice and bars are mean ± SEM. All *n* values refer to the number of mice used. *P*-values were generated for (**a, b**) by unpaired, two-tailed *t* tests, (**d**) by one-way ANOVA with Holm-Sidak’s multiple comparisons test.

**Extended Data Fig. 2:**
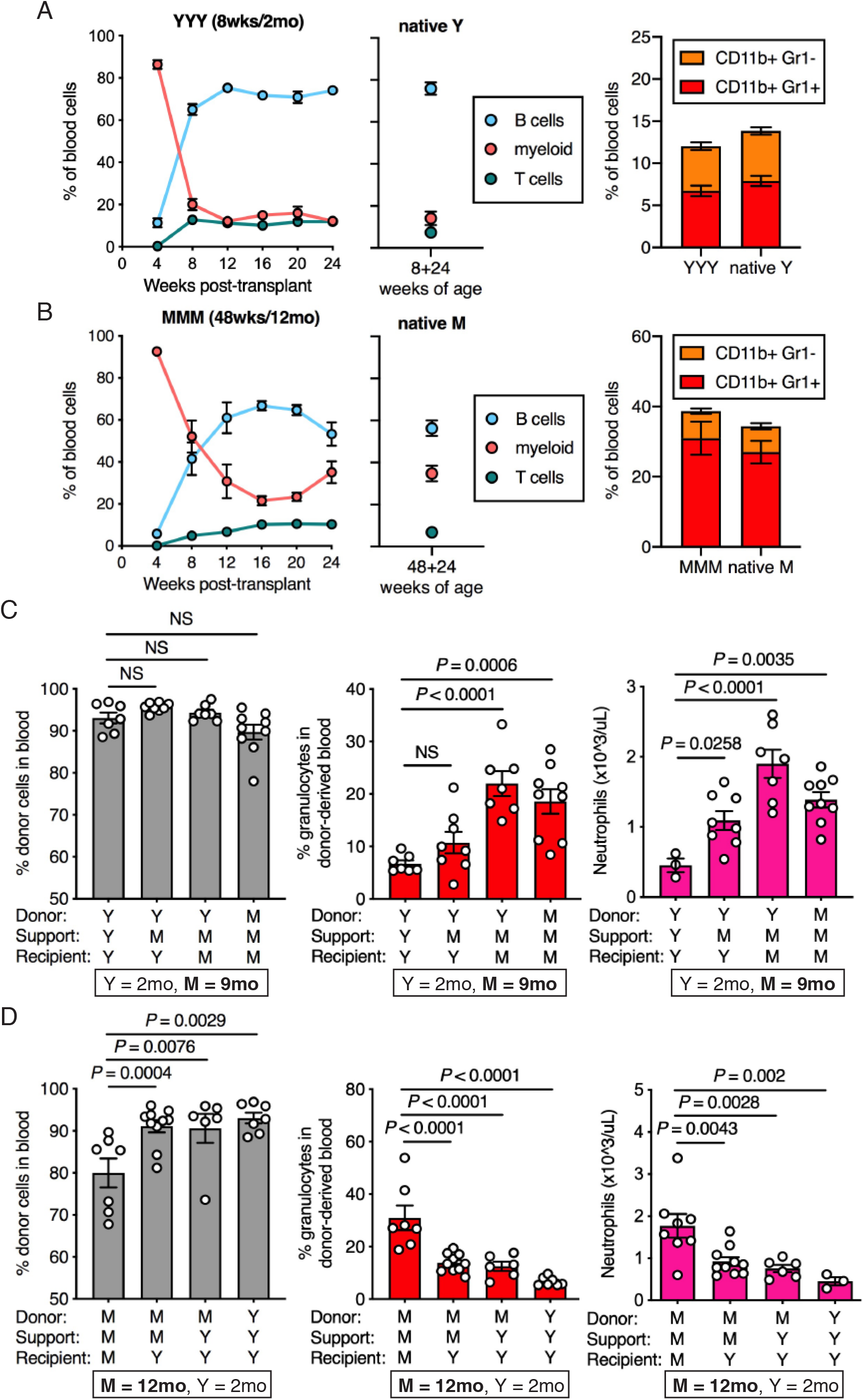
Myeloid-biased hematopoiesis at middle age is dependent on the BM microenvironment. (**a**, **b**) Left: Frequency of B cells (B220+), myeloid (CD11b+) cells, and T cells (CD3+) in peripheral blood of young (8wks;2mo) or middle-aged (48wks;12mo) control transplants over time (*n* > 5 for YYY and *n* = 7 for MMM). Center: Frequency of B, myeloid and T cells in native, non-transplanted mice age-matched to each transplant condition at the 24wks post-transplant time point (*n* = 10 for Y, *n* = 9 for M). Right: Frequency of myeloid cell subsets in peripheral blood of young or middle-aged control transplants and native, non-transplanted mice age-matched to each transplant condition (*n* = 7 for YYY, *n* = 10 for native Y, *n* = 7 for MMM, *n* = 9 for native M). (**c, d**) Left: Frequency of donor-derived cells (CD45.2+) in the blood. Center: Frequency of donor-derived granulocytes (CD11b+ Ly6c+ Ly6g+). Right: Neutrophil count. All data was obtained at 24wks post-transplant. (**a, b**) Dots (left panels) and bars (right panel) are mean ± SEM. (**c**, **d**) Dots represent individual mice and bars are mean ± SEM. All *n* values refer to the number of mice used. *P*-values were generated for (**c, d**) by one-way ANOVA with Holm-Sidak’s multiple comparisons test.

**Extended Data Fig. 3:**
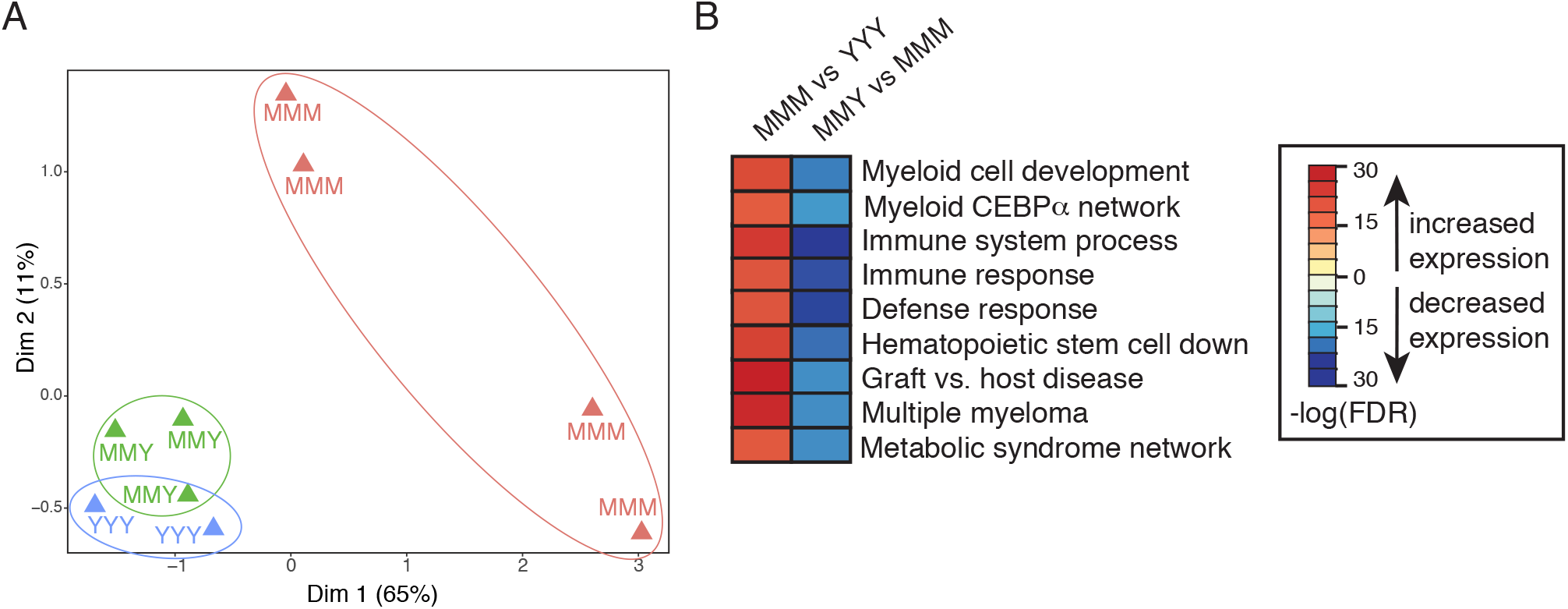
Myeloid-biased transcription signature in middle-aged LT-HSCs is rescued by a young BM microenvironment. (**a**) Multidimensional scaling (MDS) plot of 102 overlapping DEGs identified by RNA-seq data derived from MMM, MMY and YYY conditions (*n* = 4, 3, 2, from left to right). (**b**) Heatmap of gene set enrichment analysis of middle-aged vs. young LT-HSC transplant controls, and middle-aged LT-HSCs transplanted into young vs. middle-aged recipients.

**Extended Data Fig. 4:**
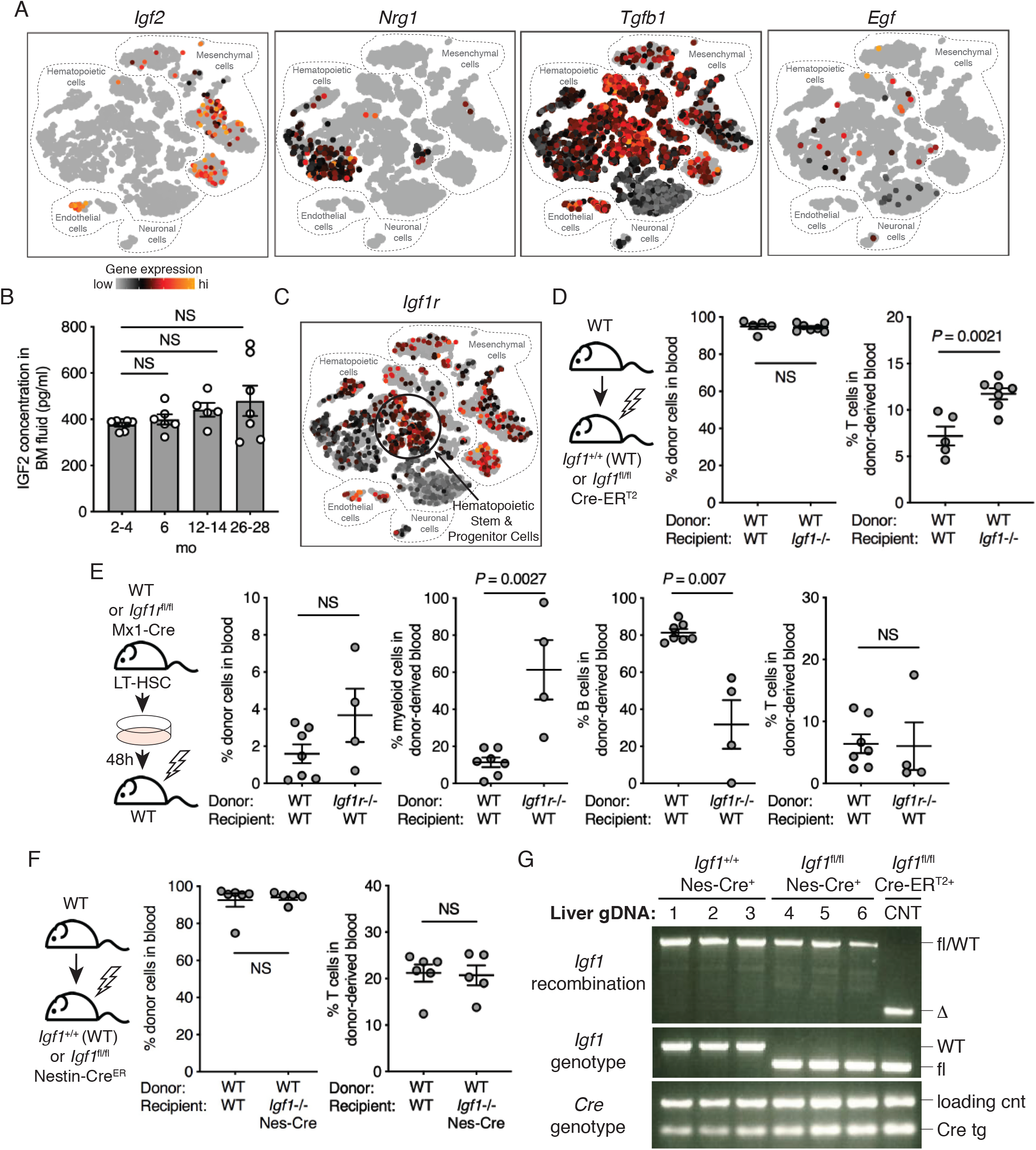
IGF1 reduction in the BM microenvironment causes myeloid-biased hematopoiesis. **a**) Expression pattern of *Igf2*, *Nrg1*, *Tgfb1*, and *Egf* in BM subsets assessed by scRNA-seq^24^. For detailed cell type annotation refer to: https://nicheview.shiny.embl.de. (**b**) IGF2 concentration in BM fluid of a single mouse femur from 2-4mo (*n* = 7), 6mo (n = 6), 12-14mo (*n* = 5) and 26-28mo (*n* = 7) mice. (**c**) Expression of *Igf1r* in BM cells by scRNA-seq^24^. (**d**) Left: Experimental design. Right: Frequency of donor cells (CD45.1+) and donor-derived T cells (CD45.1+ CD3+) in the blood at 24wks post-transplant of WT BM into lethally irradiated WT or *Igf1*^fl/fl^; CreER^T2^ recipient mice (*n* = 5, 7, from left to right). (**e**) Left: Experimental design. Right: Frequency of donor cells (CD45.2+), and donor-derived myeloid cells (CD11b+), B cells (B220+), and T cells (CD3+) in the blood at 24wks post-transplant of WT or *Igf1r*^fl/fl^ *Mx1*-Cre BM into lethally irradiated WT recipient mice (*n* = 7, 4, from left to right). (**f**) Left: Experimental design. Right: Frequency of donor cells (CD45.1+) and donor-derived T cells (CD45.1+ CD3+) in the blood at 20wks post-transplant of WT BM into lethally irradiated WT or *Igf1*^fl/fl^; Nestin-Cre^ER^ recipient mice (*n* = 6, 5, from left to right). (**g**) Top panel: *Igf1* recombination PCR, middle panel: *Igf1* genotyping PCR and bottom panel: Cre genotyping PCR. Input for all reactions was gDNA isolated from livers of tamoxifen-treated *Igf1*^+/+^ or *Igf1*^fl/fl^; Nestin-Cre^ER^ mice or recombined control (CNT; *Igf1*^fl/fl^; Cre-ER^T2^) (*n* = 3, 3, from left to right). (**b, d-f**) Dots represent individual mice and bars are mean ± SEM. *P*-values were generated for (**b**) by one-way ANOVA with Holm-Sidak’s multiple comparisons test, (**d**-**f**) by unpaired, two-tailed *t* tests.

**Extended Data Fig. 5:**
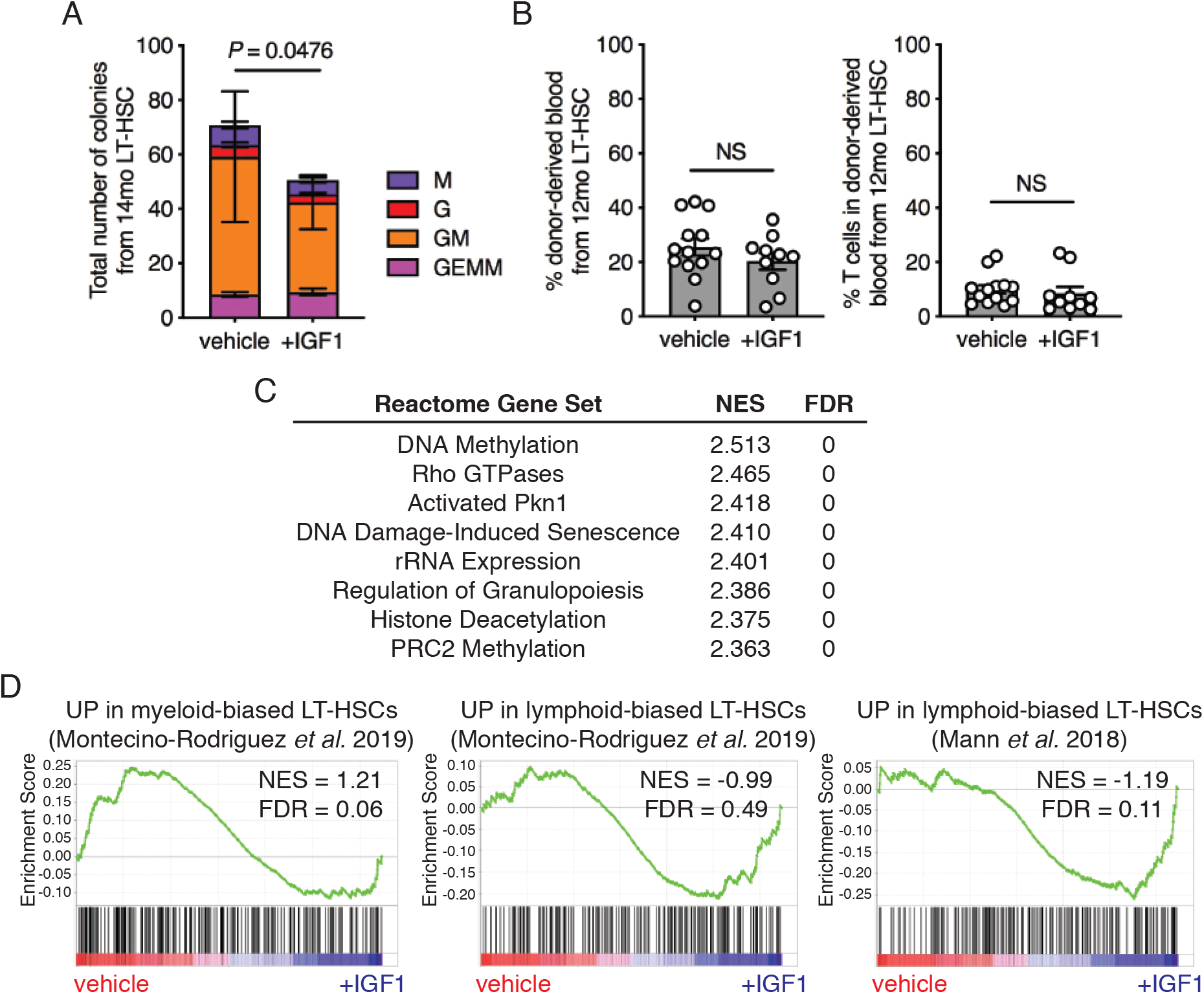
IGF1 rejuvenates middle-age LT-HSCs. (**a**) Total number of colonies derived from 14mo LT-HSCs stimulated with IGF1 or vehicle (*n* = 9). M macrophage), G (granulocyte), GM (granulocyte-macrophage), GEMM (mixed granulocyte-erythroid-macrophage-megakaryocyte). (**b**) Frequency of donor-derived cells (CD45.2+) and donor-derived T cells (CD45.2+ CD3e+) in the blood of recipient mice 8wks post-transplant of 12mo LT-HSC stimulated with vehicle (*n* = 13) or IGF1 (*n* = 10) for 7d *ex vivo*. (**c**) List of top Reactome pathways enriched in RNA-seq data of 14mo LT-HSCs stimulated with IGF1 vs. vehicle. (**d**) Left: Enrichment of myeloid-biased LT-HSC signature in vehicle-vs. IGF1-treated 14mo LT-HSCs. Center, Right: Enrichment of lymphoid-biased LT-HSC signatures in IGF1-vs. vehicle-treated 14mo LT-HSCs. (**a**, **b**) Dots represent individual mice and bars are mean ± SEM. All *n* values refer to the number of mice used. *P*-values were generated for (**a**) by paired two-tailed *t*-test and (**b**) by unpaired two-tailed *t*-test.

## Supplementary Materials

**Supplementary Table 1. Overlap of differentially expressed genes in mouse MvsY LT-HSCs, mouse OvsY LT-HSCS, and human OvsY HSCs.**

**Supplementary Table 2. Overlap of differentially expressed genes in MMMvsYYY LT-HSCs and MMYvsMMM LT-HSCs.**

**Supplementary Table 3. IPA analysis of differentially expressed genes in mouse MvsY LT-HSCs and mouse OvsY LT-HSCS.**

**Supplementary Table 4. Overlap of differentially expressed genes in MvsY vehicle-treated LT-HSCs and M+IGF1vsM+vehicle LT-HSCs.**

**Supplementary Table 5. Primer sequences for genotyping and recombination PCR.**

## Notes

### Competing Interest Statement

The authors have declared no competing interest.

## References

1 Sun, D. et al. Epigenomic profiling of young and aged HSCs reveals concerted changes during aging that reinforce self-renewal. Cell Stem Cell 14, 673–688, doi:10.1016/j.stem.2014.03.002 (2014).

2 Adelman, E. R. et al. Aging Human Hematopoietic Stem Cells Manifest Profound Epigenetic Reprogramming of Enhancers That May Predispose to Leukemia. Cancer discovery 9, 1080–1101, doi:10.1158/2159-8290.CD-18-1474 (2019).

3 Geiger, H., de Haan, G. & Florian, M. C. The ageing haematopoietic stem cell compartment. Nat Rev Immunol 13, 376–389, doi:10.1038/nri3433 (2013).

4 Mann, M. et al. Heterogeneous Responses of Hematopoietic Stem Cells to Inflammatory Stimuli Are Altered with Age. Cell Rep 25, 2992–3005 e2995, doi:10.1016/j.celrep.2018.11.056 (2018).

5 Dykstra, B., Olthof, S., Schreuder, J., Ritsema, M. & de Haan, G. Clonal analysis reveals multiple functional defects of aged murine hematopoietic stem cells. The Journal of experimental medicine 208, 2691–2703, doi:10.1084/jem.20111490 (2011).

6 Grover, A. et al. Single-cell RNA sequencing reveals molecular and functional platelet bias of aged haematopoietic stem cells. Nat Commun 7, 11075, doi:10.1038/ncomms11075 (2016).

7 Sudo, K., Ema, H., Morita, Y. & Nakauchi, H. Age-associated characteristics of murine hematopoietic stem cells. The Journal of experimental medicine 192, 1273–1280 (2000).

8 Pang, W. W. et al. Human bone marrow hematopoietic stem cells are increased in frequency and myeloid-biased with age. Proceedings of the National Academy of Sciences of the United States of America 108, 20012–20017, doi:10.1073/pnas.1116110108 (2011).

9 Flurkey, K., Currer, J. M. & Harrison, D. E. The Mouse in Aging Research. 2nd edn, 637–672 (American College Laboratory Animal Medicine (Elsevier), 2007).

10 Morrison, S. J., Wandycz, A.M., Akashi, K., Globerson, A. & Weissman, I. L. The aging of hematopoietic stem cells. Nat Med 2, 1011–16 (1996).

11 Cao, X. et al. Irradiation induces bone injury by damaging bone marrow microenvironment for stem cells. Proceedings of the National Academy of Sciences of the United States of America 108, 1609–1614, doi:10.1073/pnas.1015350108 (2011).

12 Rossi, D. J. et al. Deficiencies in DNA damage repair limit the function of haematopoietic stem cells with age. Nature 447, 725–729, doi:10.1038/nature05862 (2007).

13 Florian, M. C. et al. Cdc42 activity regulates hematopoietic stem cell aging and rejuvenation. Cell Stem Cell 10, 520–530, doi:10.1016/j.stem.2012.04.007 (2012).

14 Dulken, B. & Brunet, A. Stem Cell Aging and Sex: Are We Missing Something? Cell Stem Cell 16, 588–590, doi:10.1016/j.stem.2015.05.006 (2015).

15 Montecino-Rodriguez, E. et al. Lymphoid-Biased Hematopoietic Stem Cells Are Maintained with Age and Efficiently Generate Lymphoid Progeny. Stem Cell Reports 12, 584–596, doi:10.1016/j.stemcr.2019.01.016 (2019).

16 Ergen, A. V., Boles, N.C. & Goodell, M. A. Rantes/Ccl5 influences hematopoietic stem cell subtypes and causes myeloid skewing. Blood 119, 2500–2509, doi:10.1182/blood-2011-11-391730 (2012).

17 Maryanovich, M. et al. Adrenergic nerve degeneration in bone marrow drives aging of the hematopoietic stem cell niche. Nat Med 24, 782–791, doi:10.1038/s41591-018-0030-x (2018).

18 Kusumbe, A. P. et al. Age-dependent modulation of vascular niches for haematopoietic stem cells. Nature 532, 380–384, doi:10.1038/nature17638 (2016).

19 Verovskaya, E. V., Dellorusso, P.V. & Passegue, E. Losing Sense of Self and Surroundings: Hematopoietic Stem Cell Aging and Leukemic Transformation. Trends Mol Med 25, 494–515, doi:10.1016/j.molmed.2019.04.006 (2019).

20 de Haan, G. & Lazare, S. S. Aging of hematopoietic stem cells. Blood 131, 479–487, doi:10.1182/blood-2017-06-746412 (2018).

21 Guidi, N. et al. Osteopontin attenuates aging-associated phenotypes of hematopoietic stem cells. Embo J 36, 1463, doi:10.15252/embj.201796968 (2017).

22 Flach, J. et al. Replication stress is a potent driver of functional decline in ageing haematopoietic stem cells. Nature 512, 198–202, doi:10.1038/nature13619 (2014).

23 Young, K. et al. Progressive alterations in multipotent hematopoietic progenitors underlie lymphoid cell loss in aging. The Journal of experimental medicine 213, 2259–2267, doi:10.1084/jem.20160168 (2016).

24 Baccin, C. et al. Combined single-cell and spatial transcriptomics reveal the molecular, cellular and spatial bone marrow niche organization. Nat Cell Biol, doi:10.1038/s41556-019-0439-6 (2019).

25 Mendez-Ferrer, S. et al. Mesenchymal and haematopoietic stem cells form a unique bone marrow niche. Nature 466, 829–834, doi:10.1038/nature09262 (2010).

26 Greenbaum, A. et al. CXCL12 in early mesenchymal progenitors is required for haematopoietic stem-cell maintenance. Nature 495, 227–230, doi:10.1038/nature11926 (2013).

27 Ding, L., Saunders, T.L., Enikolopov, G. & Morrison, S. J. Endothelial and perivascular cells maintain haematopoietic stem cells. Nature 481, 457–462, doi:10.1038/nature10783 (2012).

28 Kfoury, Y. & Scadden, D. T. Mesenchymal cell contributions to the stem cell niche. Cell Stem Cell 16, 239–253, doi:10.1016/j.stem.2015.02.019 (2015).

29 Caselli, A. et al. IGF-1-mediated osteoblastic niche expansion enhances long-term hematopoietic stem cell engraftment after murine bone marrow transplantation. Stem Cells 31, 2193–2204, doi:10.1002/stem.1463 (2013).

30 Burns, K. A. et al. Nestin-CreER mice reveal DNA synthesis by nonapoptotic neurons following cerebral ischemia hypoxia. Cereb Cortex 17, 2585–2592, doi:10.1093/cercor/bhl164 (2007).

31 Chen, K. G., Johnson, K.R. & Robey, P. G. Mouse Genetic Analysis of Bone Marrow Stem Cell Niches: Technological Pitfalls, Challenges, and Translational Considerations. Stem Cell Reports 9, 1343–1358, doi:10.1016/j.stemcr.2017.09.014 (2017).

32 Powell-Braxton, L. et al. IGF-I is required for normal embryonic growth in mice. Genes Dev 7, 2609–2617, doi:10.1101/gad.7.12b.2609 (1993).

33 Yakar, S. et al. Circulating levels of IGF-1 directly regulate bone growth and density. J Clin Invest 110, 771–781, doi:10.1172/JCI15463 (2002).

34 Baryawno, N. et al. A Cellular Taxonomy of the Bone Marrow Stroma in Homeostasis and Leukemia. Cell 177, 1915–1932 e1916, doi:10.1016/j.cell.2019.04.040 (2019).

35 Kineman, R. D., Del Rio-Moreno, M. & Sarmento-Cabral, A. 40 YEARS of IGF1: Understanding the tissue-specific roles of IGF1/IGF1R in regulating metabolism using the Cre/loxP system. J Mol Endocrinol 61, T187–T198, doi:10.1530/JME-18-0076 (2018).

36 Pietras, E. M. et al. Functionally Distinct Subsets of Lineage-Biased Multipotent Progenitors Control Blood Production in Normal and Regenerative Conditions. Cell stem cell 17, 35–46, doi:10.1016/j.stem.2015.05.003 (2015).

37 Wilkinson, A. C. et al. Long-term ex vivo haematopoietic-stem-cell expansion allows nonconditioned transplantation. Nature 571, 117–121, doi:10.1038/s41586-019-1244-x (2019).

38 Liang, Y., Van Zant, G. & Szilvassy, S. J. Effects of aging on the homing and engraftment of murine hematopoietic stem and progenitor cells. Blood 106, 1479–1487, doi:10.1182/blood-2004-11-4282 (2005).

39 Rossi, D. J. et al. Cell intrinsic alterations underlie hematopoietic stem cell aging. Proceedings of the National Academy of Sciences of the United States of America 102, 9194–9199, doi:10.1073/pnas.0503280102 (2005).

40 Dykstra, B., Olthof, S., Schreuder, J., Ritsema, M. & de Haan, G. Clonal analysis reveals multiple functional defects of aged murine hematopoietic stem cells. J Exp Med 208, 2691–2703, doi:10.1084/jem.20111490 (2011).

41 Hede, M. S. et al. E-peptides control bioavailability of IGF-1. PLoS One 7, e51152, doi:10.1371/journal.pone.0051152 (2012).

42 Cheng, C. W. et al. Prolonged fasting reduces IGF-1/PKA to promote hematopoietic-stem-cell-based regeneration and reverse immunosuppression. Cell Stem Cell 14, 810–823, doi:10.1016/j.stem.2014.04.014 (2014).

43 Tang, D. et al. Dietary restriction improves repopulation but impairs lymphoid differentiation capacity of hematopoietic stem cells in early aging. J Exp Med 213, 535–553, doi:10.1084/jem.20151100 (2016).

44 Ho, T. T. et al. Autophagy maintains the metabolism and function of young and old stem cells. Nature 543, 205–210, doi:10.1038/nature21388 (2017).

45 Subramanian, A. et al. Gene set enrichment analysis: a knowledge-based approach for interpreting genome-wide expression profiles. Proceedings of the National Academy of Sciences of the United States of America 102, 15545–15550, doi:10.1073/pnas.0506580102 (2005).

46 Boulais, P. E. et al. The Majority of CD45(-) Ter119(-) CD31(-) Bone Marrow Cell Fraction Is of Hematopoietic Origin and Contains Erythroid and Lymphoid Progenitors. Immunity 49, 627–639 e626, doi:10.1016/j.immuni.2018.08.019 (2018).

47 Huang, S. et al. An improved protocol for isolation and culture of mesenchymal stem cells from mouse bone marrow. J Orthop Translat 3, 26–33, doi:10.1016/j.jot.2014.07.005 (2015).

