## Supplementary Table 5 for "Hematopoietic Stem and Progenitor Cell Aging is Initiated at Middle Age Through Decline in Local Insulin-Like Growth Factor 1 (IGF1)"

**Supplementary Table 4. Primer sequences for genotyping and recombination PCR**

**Primer Name	Application(s)	Sequence (5’ to 3’)**

Igf1flox Mut F	genotyping	AAA CCA CAC TGC TCG ACA TTG

Igf1flox Wt F	genotyping	GGC AAA TGG AAA TCC TAT GTC T

Igf1flox R	genotyping,	CAC TAA GGA GTC TGT ATT TGG ACC

recombination

Igf1flox Mut F	recombination	AGC CTC TCA ACT AAG ACA ATA

Cre tg84	genotyping	GCG GTC TGG CAG TAA AAA CTA TC

Cre tg85	genotyping	GTG AAA CAG CAT TGC TGT CAC TT

Cre F	genotyping	 CTA GGC CAC AGA ATT GAA AGA TCT

Cre R	genotyping	GTA GGT GGA AAT TCT AGC ATC ATC C

Igf1 RT F	real-time PCR	CCGAGGGGCTTTTACTTCAAC

Igf1 RT R	real-time PCR	CAGTCTCCTCAGATCACAGCT

B2M RT F	real-time PCR	TTCTGGTGCTTGTCTCACTGA

B2M RT R	real-time PCR	CAGTATGTTCGGCTTCCCATTC
